# Actin Networking Collapse Under STAT5A Deficiency Drives Mitochondrial Dysfunction and Autocrine IFN-β Production

**DOI:** 10.1101/2025.05.26.656095

**Authors:** Boyue Sun, Mathumathi Krishnamohan, Reinat Nevo, Eyal Zoler, Benjamin Geiger, Gideon Schreiber

**Affiliations:** Department of Biomolecular Sciences, Weizmann Institute of Science, Rehovot, Israel; Department of Immunology and Regenerative Biology, Weizmann Institute of Science, Rehovot, Israel

**Keywords:** STAT5A, Actin Remodeling, ACTN1, Mitochondrial Dynamics, Type I interferon, cGAS-STING pathway, genomic instability, JAK-STAT Signaling Cascade

## Abstract

Coordination between actin cytoskeleton networking and mitochondrial organization and health underpins cellular homeostasis. Here, we found STAT5A to be a pivotal transcription factor that sustains the expression of key actin regulators, including ACTN1. STAT5A, but not STAT5B deficiency dismantles F-actin architecture leading to amorphous cell shape, reduced cellular motility, and corrals mitochondria around the nucleus. This actin networking disruption impairs mitochondrial DRP1 recruitment and dynamic equilibrium, leading to ROS-associated DNA damage and cGAS–STING mediated type I IFN production. Consequently, this establishes a chronically IFN-stimulated state in neighboring cells. Conversely, overexpression of STAT5A increases actin cytoskeleton networking and promotes faster cell motility. Moreover, ectopic expression of ACTN1 in the background of STAT5A-knockout cells is sufficient to restore the actin cytoskeleton organization and mitochondrial network morphology, eliminating DNA damage and IFN-signaling. Giving the importance of actin in cellular homeostasis, our findings place actin abundance, as regulated by the STAT5A–ACTN1 axis as essential for linking cytoskeletal integrity with mitochondrial health, restraining aberrant innate immune activation.

**Graphic Abstract:** 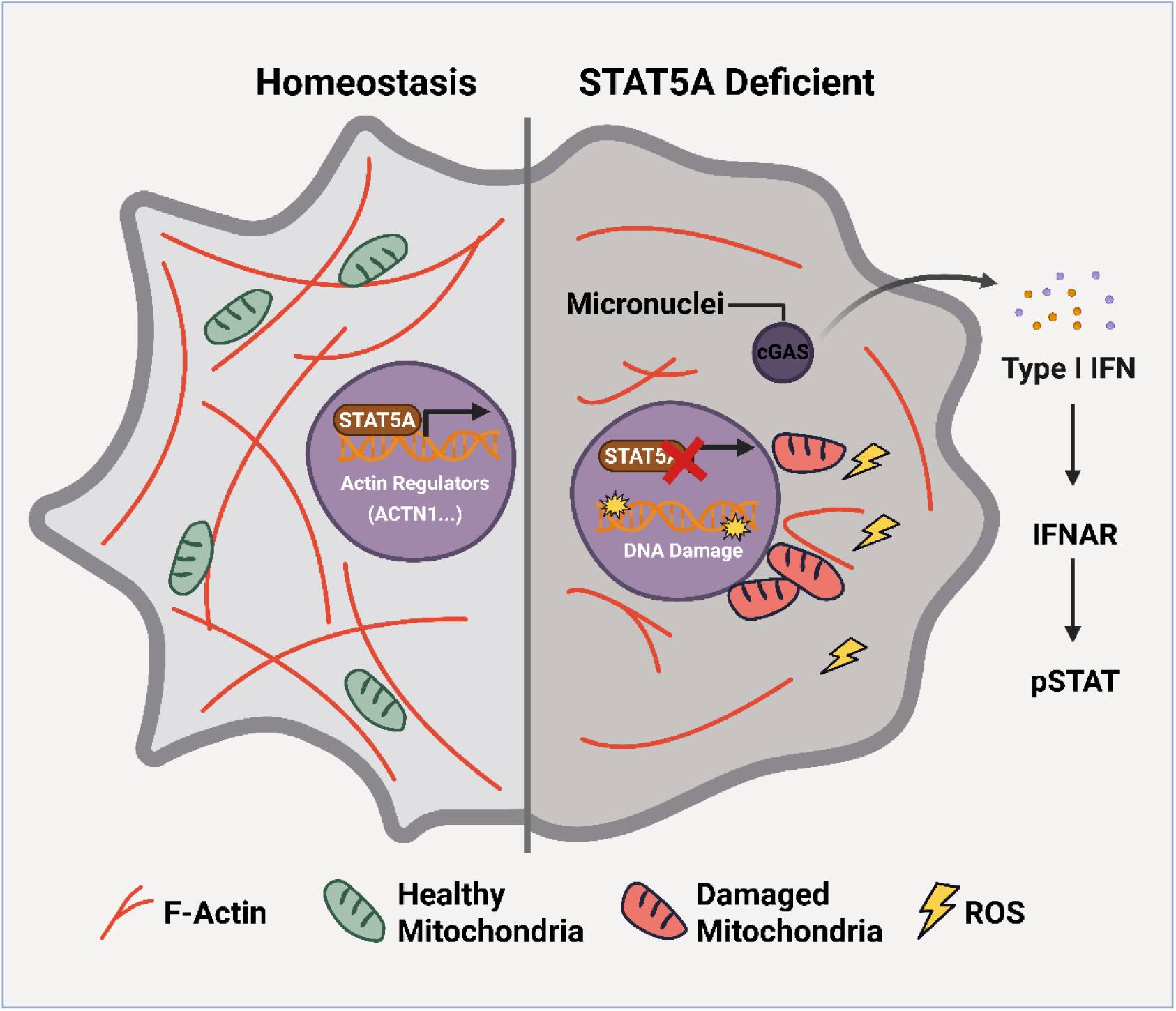

**Key points:** STAT5A deficiency disrupts cytoskeletal integrity via ACTN1 loss.

The disrupted actin-cytoskeleton impairs mitochondria organization, causing ROS production and DNA damage.

STAT5A knockout activates cGAS–STING signaling and IFN-β production.

Restoring ACTN1 rescues cytoskeleton, mitochondrial function, and immune balance.

## Introduction

Signal transducer and activator of transcription 5 (STAT5) is a critical mediator of cytokine and growth factor signaling with two highly homologous isoforms, STAT5A and STAT5B. Canonically, STAT5 proteins function as latent cytosolic transcription factors: upon tyrosine phosphorylation by JAK kinases downstream of cytokines (such as interleukins, growth hormone, prolactin, and erythropoietin), STAT5A/B dimerize and translocate to the nucleus to regulate target gene expression^1,2^. Through this pathway, STAT5 controls diverse biological processes critical for cellular development, differentiation, and proliferation^1,2^.

Dysregulation of STAT5 signaling, is implicated in the pathogenesis of various diseases, including autoimmune conditions such as rheumatoid arthritis, systemic lupus erythematosus, and multiple sclerosis, as well as diverse malignancies such as leukemia, lymphoma, breast, and prostate cancers^3–6^.Targeting STAT5 in cancer therapy is promising, with recent advancements in small molecule inhibitors that either degrade STAT5 or inhibit its phosphorylation, showing efficacy in preclinical studies ^7–9^. In addition to its canonical nuclear role, STAT5 has non-canonical functions, including organelle maintenance and cell stress protection. In human endothelial cells, knockdown of STAT5 led to rapid organelle disorganization: the ER underwent dramatic cystic dilation with aberrant accumulation of ER-shaping proteins on the cyst membranes, the Golgi complex fragmented, and the nucleus became misshapen^10^. Functionally, these structural perturbations were accompanied by impaired intracellular trafficking (reduced anterograde transport from ER to Golgi) and an unfolded protein response (increased GRP78/BiP), indicating ER stress^10^. In addition, STAT5A but not STAT5B, was specifically found to protect cells from various stress conditions, including oxidative stress and hypoxia, helping in cell survival and adaptation to stress^11–14^.

Actin filament networks, a principal component of the cytoskeleton, have emerged as critical regulators of organelle form and function, including mitochondrial organization and health^15–18^. Actin networks contribute significantly to mitochondrial positioning by mediating both transport and anchoring mechanisms. For example, Myosin-19 binds to mitochondria and propels them along actin filaments toward the cell periphery, whereas polymerizing actin networks can encapsulate mitochondria in cage-like structures, limiting their motility^15–18^. The actin assembly is also a co-factor for the canonical fission machinery, illustrating a non-muscle contractile force that helps divide mitochondria. Actin scaffolds help recruit and potentiate the GTPase DRP1, the main executor of mitochondrial fission, leading to efficient scission of the organelle^15–18^. Beyond morphology and placement, actin impacts mitochondrial function during cellular stress. Recent work demonstrates that when mitochondria become dysfunctional, F-actin rapidly assembles around the damaged mitochondria within, mechanically preventing pathological swelling or fragmentation and facilitating a metabolic shift toward glycolysis^18^.

Proper mitochondrial fission and clearance, which depend on actin networks, are necessary to isolate and remove ROS-generating mitochondria. If actin dynamics are disrupted, cells often experience an increase in ROS levels that inflict damage on other cellular components, including nuclear DNA^19^. Oxidatively damaged mtDNA leaking through pores in the mitochondria, or genomic DNA then reach the cytosol in the form of micronuclei that is sensed as foreign by cGAS. Once triggered, the cGAS– STING axis induces an antiviral-like state, with robust production of IFN-β and IFN-stimulated genes that amplify inflammatory cascades^20^. This innate DNA-sensing pathway is crucial for host defense and for clearance of abnormal cells – for example, cGAS–STING can detect tumor-derived DNA or mitochondrial DNA from damaged cells and promote anti-tumor immunity or cell death. On the other hand, inappropriate or chronic activation of cGAS–STING has been implicated in a spectrum of disorders, and is a potential therapeutic target in infectious, autoimmune, and degenerative diseases as well as cancer^19^. In autoimmune syndromes like systemic lupus erythematosus, failure to clear nuclear or mitochondrial DNA debris leads to persistent cGAS activation and pathological IFN production. In inflammatory heart and lung diseases, mtDNA release during cell injury can drive STING-mediated tissue damage. Even in the brain, cGAS–STING emerged as a key driver of “sterile” neuroinflammation that can induce a pro-inflammatory state implicated in neurodegenerative diseases^19^. In this context, mitochondrial maintenance and cytoskeletal integrity are highly relevant to the cell’s immunological quietude.

In this study, we observed that STAT5A, traditionally known as a cytokine-activated transcription factor, functions as a regulator that maintains actin cytoskeletal networks support for mitochondrial function and prevents cells from inadvertently initiating type I IFN responses. The concept that a STAT protein coordinately regulates actin organization, mitochondrial homeostasis, and innate immune quiescence is a novel paradigm that bridges signaling pathways normally considered distinct. The findings could reveal a novel facet of cellular homeostasis and suggest innovative strategies for therapeutic intervention in STAT5-driven cancers or inflammatory diseases where mitochondria and innate immunity intersect.

## Results

### Altered IFN responses in STAT KO HeLa Cells

To dissect the individual contributions of STAT family members to cellular signaling, we generated a comprehensive panel of CRISPR-Cas9–engineered STAT-knockout HeLa cell lines. Loss of STAT1, STAT2, STAT3, STAT4, STAT5A, STAT5B and STAT6 were confirmed by Sanger sequencing and by the absence of the corresponding proteins on Western blots (Figure 1A and Figure S1A). Global transcriptional consequences of each KO were assessed by RNA-seq performed under basal conditions and after 16 h of IFN-β stimulation. Principal-component analysis (PCA) of these datasets (Figure 1B) revealed clear separation of the transcriptional profiles: the first component (PC1; 65.6 % of the total variance) isolated IFN-β-treated STAT5A KO samples from all other groups. To visualize the interferon-stimulated gene (ISG) response in greater detail, ISGs were stratified into high, moderate and low induction groups according to their log2 fold-change values (Figure 1C). STAT5A deficiency produced broad hyper-induction across all three bins, whereas STAT6 loss selectively attenuated ISG induction, most markedly within the moderate- and low-induction categories. Repression of genes normally down-regulated by IFN-β were unaffected in STAT5A KO cells but partially relieved in STAT6 KO cells. Moreover, STAT5A KO cells were more sensitive to IFN-β stimulation, as evidenced by increased ISG expression at both low and high concentrations of IFN-β treatment. Intriguingly, even without treatment, STAT5A KO cells exhibited an elevated basal level of MX1 and IDO1 expression (Figure 1D), demonstrating that the homeostatic shift elicited by STAT5A deficiency.

**Figure 1.**
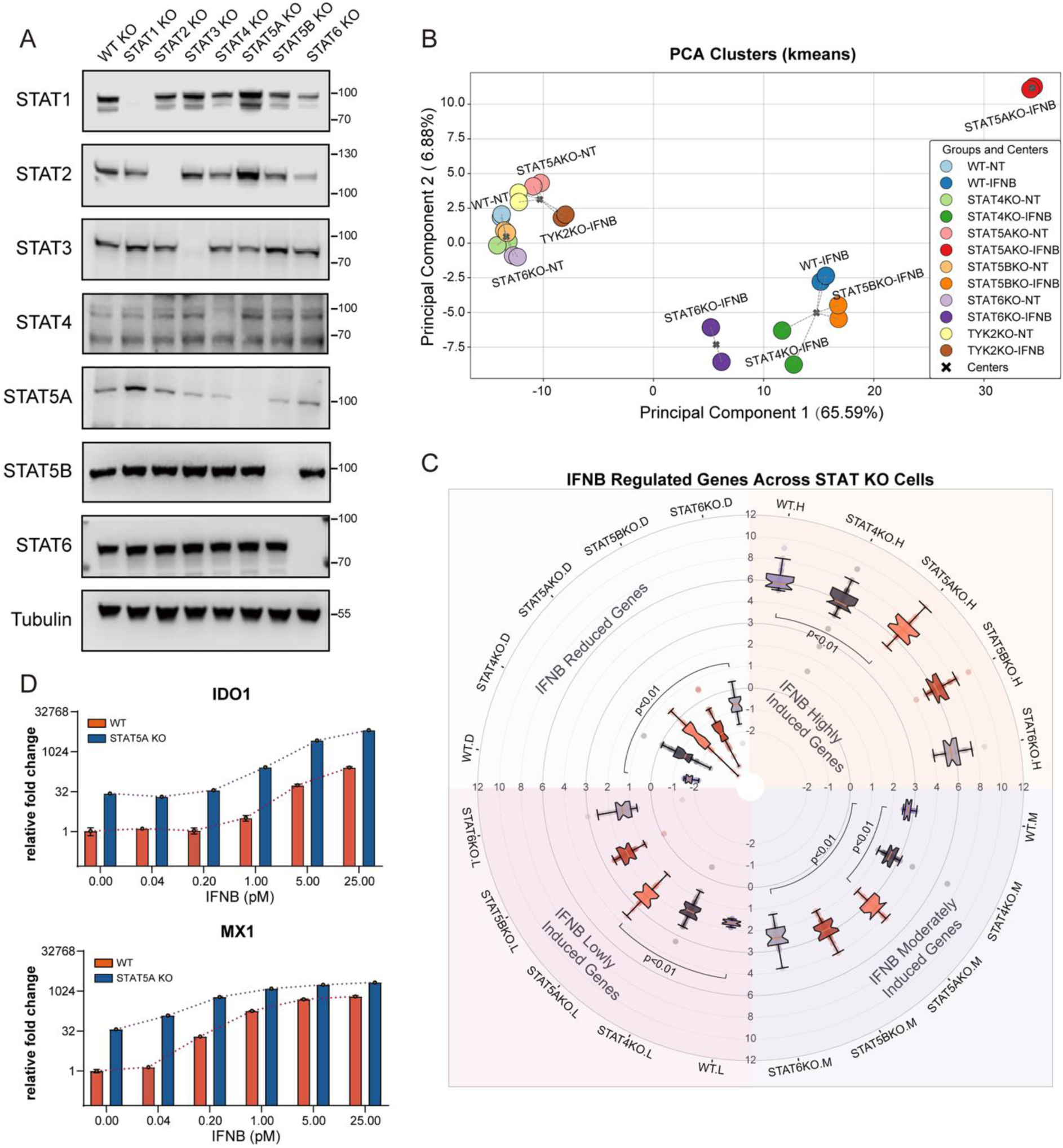
STAT5A KO Enhances IFN-β Response. (A) Western blot analysis of WT, STAT1 KO, STAT2 KO, STAT3 KO, STAT4 KO, STAT5A KO, STAT5B KO, and STAT6 KO cell lines were probed with specific antibodies. (**B**) Principal Component Analysis with K-means clustering of ISG expression profiles. The plot shows the first two principal components of ISG expression profiles across WT and mutant HeLa cells, with and without IFN-β treatment, accounting for 65.59% and 6.88% of the variance. Five distinct clusters were identified using K-means clustering. Each point represents an individual sample, colored according to genotype and treatment; cluster centers (indicated with an ‘×’) highlight the groupings. (**C**) Response of WT and STAT5A KO cells to increasing concentrations of IFN-β. WT and STAT5A KO HeLa cells were treated with IFN-β for 16 hours, and the expression levels of MX1 and IDO1 were quantified by qPCR. (D) Radial plot displaying differential expression of IFN-β-induced genes. Genes are categorized based on their induction levels into IFN-β-highly induced, moderately induced, lowly induced or reduced groups. Each quadrant represents one of these categories, with boxplots showing the distribution of gene expression for each condition across WT and mutant HeLa cells.

### STAT5A KO Leads to Constitutive Activation of the IFNAR-JAK-STAT Pathway via Autocrine Type I IFN Production

The elevated basal ISG expression observed in STAT5A KO cells suggests the involvement of JAK-STAT pathway activation. To investigate this possibility, we performed Western blot analysis on untreated cells, revealing increased phosphorylation of STAT1 and STAT2 specifically in STAT5A KO cells (Figure 2A). To further confirm the direct role of STAT5A in this phenotype, we reintroduced STAT5A into STAT5A KO cells through stable transfection. Restoration of STAT5A expression abolished STAT1 and STAT2 phosphorylation as well as the basal ISG expression, clearly demonstrating that the observed activation was directly attributable to the absence of STAT5A (Figure 2B and S2A).

**Figure 2.**
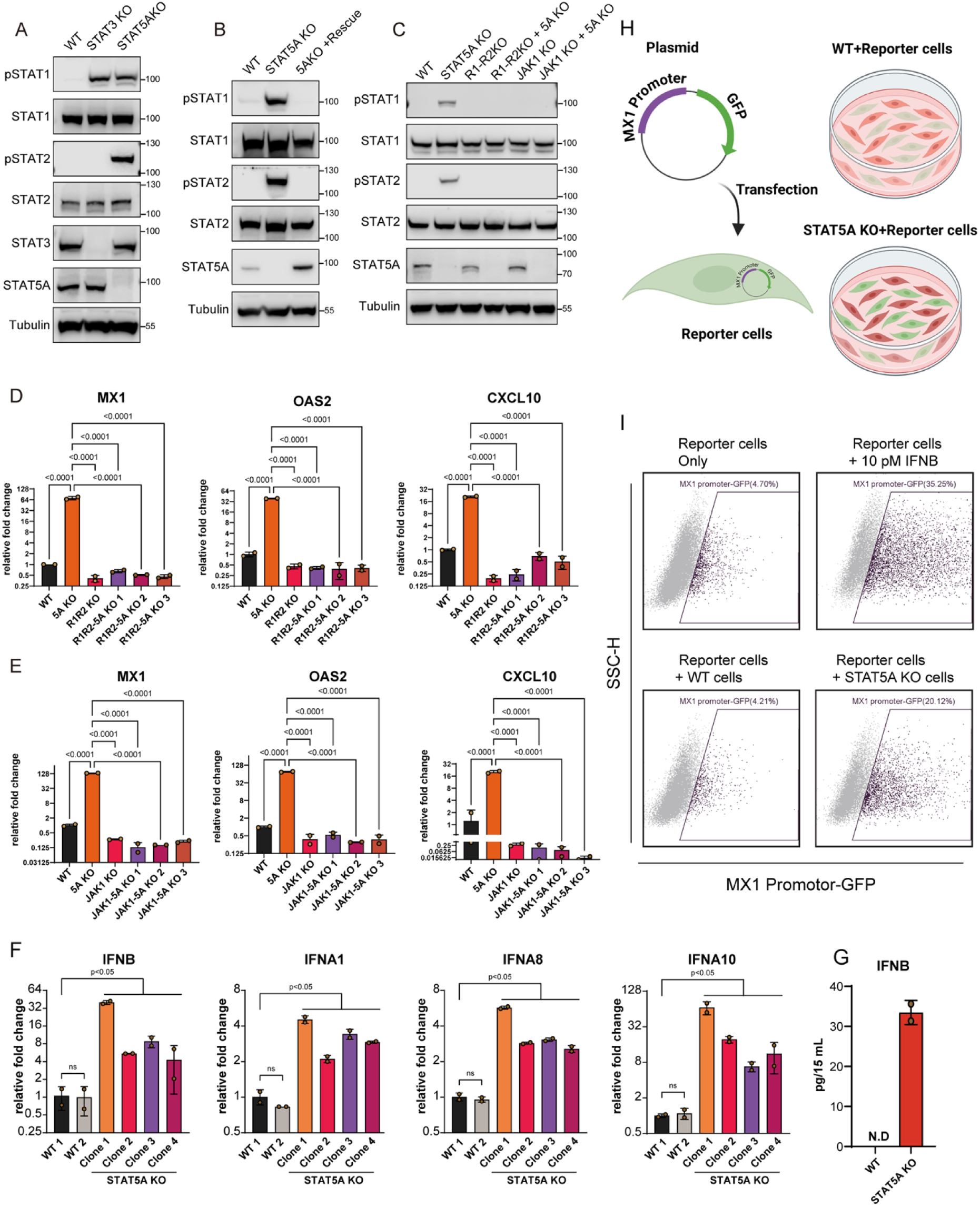
STAT5A KO Leads to Constitutive Activation of the IFNAR-JAK-STAT Pathway via Autocrine Type I IFN Production. (A-C) Western blot analysis of STAT levels without external type I IFN addition. (A) STAT1, 2, 3 and 5 levels in WT, STAT3 KO, and STAT5A KO HeLa cells. (B) STAT1, 2 and 5 in WT, STAT5A KO, and STAT5A rescue cells. (C) Analysis of pSTAT1, 2 and 5 levels in WT and KO cells. R1 and R2 represent IFNAR1 and IFNAR2, respectively, while 5A represents STAT5A. (D-F) qPCR analysis of ISG and IFN expression. (D) Relative expression of ISGs in WT, STAT5A KO, IFNAR1/2 KO, and IFNAR1/2-STAT5A triple KO clones. (E) Relative expression of ISGs in WT, STAT5A KO, JAK1 KO, and JAK1-STAT5A double KO clones. (F) Relative expression of type I IFN genes in WT and STAT5A KO HeLa cells. No external type I IFN was added in D-F. (G) Quantification of IFN-β protein in cell culture supernatants in WT and STAT5A KO cells. “N.D.” indicates that IFN-β levels were below the detection limit in WT cells. (H-I) Validation of type I IFN production by STAT5A KO cells using an MX1 promoter-GFP reporter. (H) Schematic representation of the experimental setup: WT cells transfected with an MX1 promoter-GFP plasmid were co-cultured with either WT or STAT5A KO cells. (I) Flow cytometry analysis of GFP expression in different experimental conditions. 10 pM IFN-β treatment of 12 hours was used as a positive control. For Figure 2D-2F, statistical significance was calculated using one-way ANOVA followed by Dunnett’s post hoc test. Error bars indicate standard deviation. Data shown are representative of at least three independent experiments.

To delineate the involvement of upstream signaling components, we generated IFNAR1-IFNAR2-STAT5A triple KO cells and JAK1-STAT5A double KO cells. Western blot analysis showed that cells lacking STAT5A in combination with either JAK1 or IFNARs did not exhibit increased phosphorylation of STAT1 and STAT2. This finding underscores the necessity of IFNAR and JAK1 signaling for the STAT5A KO-induced phenotype (Figure 2C). Consistent with these results, quantitative PCR (qPCR) analysis confirmed that ISG expression induced by STAT5A KO depends on functional IFNAR receptor complexes and JAK1 signaling pathways (Figures 2D and 2E).

The dependency of IFNAR and JAK1 signaling for STAT phosphorylation and ISG expression raised the hypothesis that type I IFNs might be produced in STAT5A KO cells. To test this hypothesis, we performed qPCR analysis, which revealed significantly elevated expression levels of type I IFN genes in multiple STAT5A KO clones compared to WT cells (Figure 2F). Quantification of IFN-β protein in the cell culture media using the LEGENDplex assay indicated that IFN-β was undetectable in WT cells but was present at approximately 30 pg/15 mL in STAT5A KO cell media, confirming active secretion of IFN-β protein (Figure 2G). To validate the biological activity of secreted IFN-β, we utilized a reporter assay employing an MX1 promoter-driven GFP construct, which is highly sensitive to even low levels of type I IFNs. HeLa-WT cells transfected with this reporter construct were co-cultured with either WT or STAT5A KO cells. Reporter cells co-cultured with STAT5A KO cells exhibited approximately 20% GFP-positive cells, compared to around 35% positivity induced by treatment with 10 pM IFN-β. These results clearly demonstrate activation of the MX1 promoter by IFN-β secreted from STAT5A KO cells (Figures 2H and 2I). Taken together, these findings indicate that the knockout of STAT5A induces the production and secretion of biologically active type I IFNs, resulting in constitutive activation of the IFNAR-JAK-STAT signaling pathway.

### STAT5A Knockout alters Gene Expression of Key Cellular Components

The constitutive activation of IFNAR signaling and autocrine secretion of Type I IFNs prompted us to investigate the underlying causes of these phenomena. Transcriptomic analysis revealed significant differential expression in STAT5A KO cells compared to WT cells (or STAT5B KO cells), with 279 genes upregulated (Log2FC > 1, p < 0.05) and 353 genes downregulated (Log2FC < -1, p < 0.05) (Figure 3A). Gene set enrichment analysis (GSEA) of these differentially expressed genes demonstrated a pronounced upregulation of IFN-related and immune response pathways, similar to cells treated with type I IFNs. Notably, the “Interferon Alpha/Beta Signaling” pathway displayed the highest positive z-score (4.583), encompassing 29.6% of the analyzed gene set, underscoring a robust type I IFN response. Additionally, inhibition of the “Coronavirus Replication Pathway” and activation of the “ISGylation Signaling Pathway” further emphasized the heightened immune activation in STAT5A KO cells (Figure 3B). These findings align with the results presented in Figure 2. Further analysis utilizing correlation network analysis via the Ingenuity Pathway Analysis (IPA) platform, based on all differentially expressed genes, provided deeper insights into the molecular consequences of STAT5A knockout (Figure 3C). Correlation network indicated that STAT5A KO cells exhibit molecular profiles analogous to cells treated with type I IFNs, characterized by inhibition of the viral life cycle. Pathways associated with apoptosis and necrosis were also activated, consistent with established roles of interferons in mediating cell death processes. Conversely, cellular and tissue development pathways were predominantly suppressed, consistent with previous in vivo and in vitro studies highlighting developmental impairments associated with STAT5A deficiency^2,5,21^. Furthermore, STAT5A deficiency impacted filament and cytoskeletal formation as well as cancer cell migration and invasion (Figure 3C). Detailed transcriptomic analysis specifically identified reduced expression levels of genes critical for actin cross-linking and regulation, such as ACTN1, FLNA, and MYH9, among others (Figure 3D). These transcriptional changes were validated by qPCR, confirming the decreased expression of these structural components (Figure 3E and S3D).

**Figure 3.**
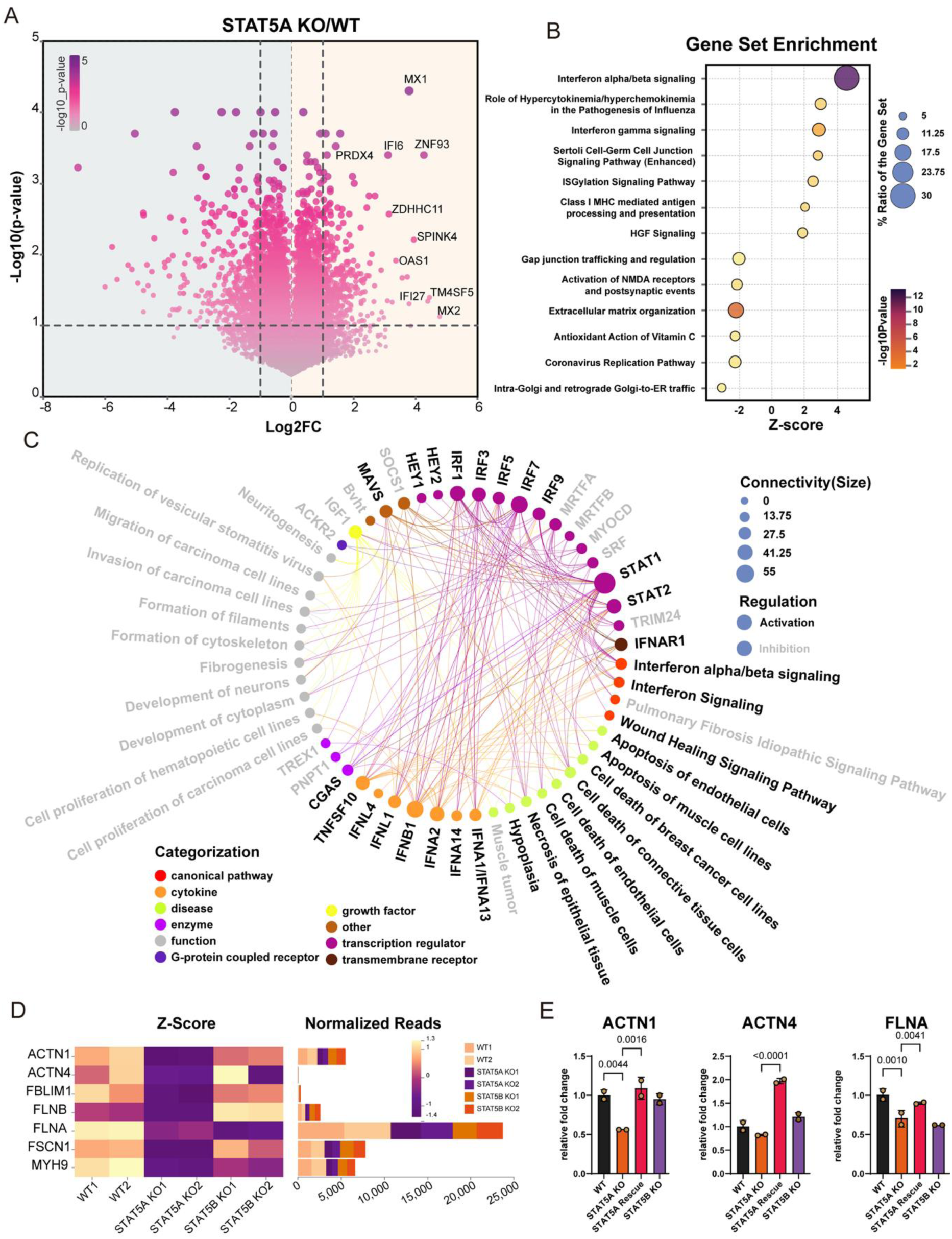
STAT5A Knockout alters Gene Expression of Key Cellular Components. (A) Volcano plot shows the expression levels of genes in STAT5A KO cells compared to WT controls. Dashed lines indicate thresholds for statistical significance (p < 0.05) and fold-change (log2FC > 1 or < -1). (B) GSEA of differentially expressed genes. The plot highlights the enriched biological pathways, with the x-axis representing the Z-score and the y-axis listing the enriched pathways. The size of each circle corresponds to the percentage ratio of the gene set, while the color gradient reflects the -log10 p-value of enrichment. (C) Correlation network map of key genes and pathways generated using the IPA platform on the RNA-seq dataset. The size of each node reflects the connectivity or interaction strength, with larger nodes indicating a higher number of interactions. Black labels are for activated processes, while grey labels indicate inhibited processes. (D) Heatmaps displaying RNA-seq analysis of actin regulating genes in WT, STAT5A KO, and STAT5A rescue cells. Z-score heatmaps represent differential gene expressions, while bar plots illustrate normalized read counts. (E) qPCR analysis of cytoskeleton and adhesion genes expression in WT, STAT5A KO, STAT5A Rescue cells. For Figure 3E statistical significance was calculated using one-way ANOVA followed by Dunnett’s post hoc test. Error bars indicate standard deviation. Data shown are representative of at least three independent experiments.

### STAT5A Knockout Leads to disruption of the Actin network

The transcriptomic data (Figures 3D and 3E) let us to image STAT5A deficient cells under a confocal microscope. These images revealed that STAT5A deficiency significantly disrupts the actin cytoskeletal architecture. In WT cells, actin filaments are evenly distributed throughout the basal region, characterized by well-organized longitudinal and peripheral stress fibers aligned along the cell cortex, supporting a structured and polarized morphology. In contrast, STAT5A KO cells exhibit a flattened and irregular morphology with markedly disorganized actin filaments and significantly reduced central filamentous structures. Peripheral stress fibers are also diminished, resulting in compromised integrity of the actin network. Notably, re-expression of STAT5A in deficient cells successfully restored normal actin filament architecture (Figure 4A). Similar cytoskeletal defects were consistently observed in fixed, Mito-TEMPO-treated cells stained with phalloidin, ruling out potential artifacts from LifeAct expression or oxidative cell stress (Figure 4B). Additionally, STAT5A KO cells demonstrated clear alterations in cell size (Figure 4J). Comparable cytoskeletal abnormalities were also identified in STAT5A KO B16F1 cells (Figure S3A), emphasizing the critical and specific role of STAT5A in maintaining actin cytoskeletal network integrity.

**Figure 4.**
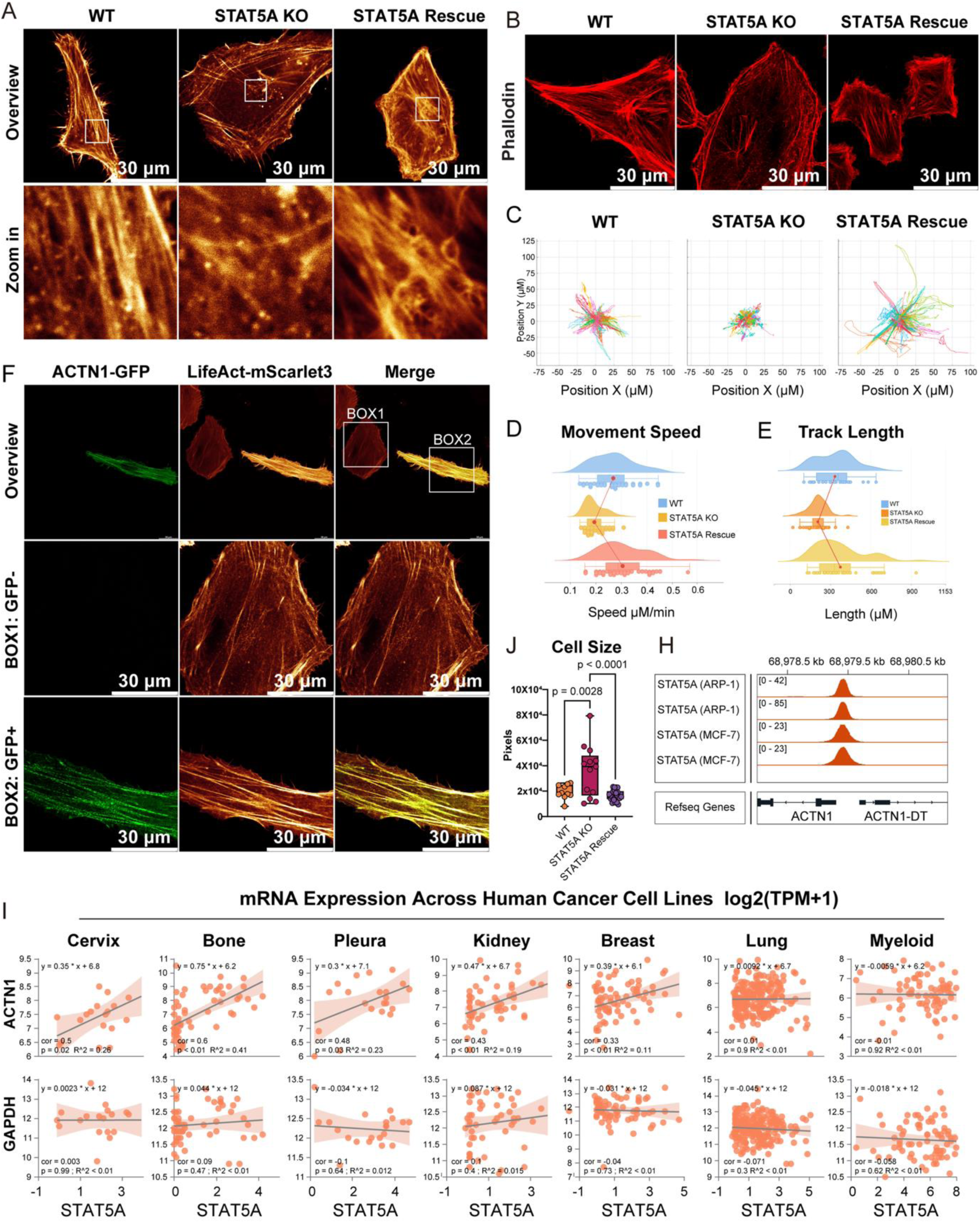
STAT5A Knockout Leads to disruption of the Actin network. (A) Confocal microscopy images at the basal plane of WT, STAT5A KO, and STAT5A-rescued cells overexpressing LifeAct-mScarlet3. (B) Confocal microscopy images at the basal plane of WT, STAT5A KO, and STAT5A-rescued cells treated with 10 µM Mito-TEMPO and stained for actin filaments using phalloidin (red). (C-E) Cell Migration Trajectories and Speed Analysis. (C) Representative migration trajectories of WT, STAT5A KO, and STAT5A rescue cells recorded over 273 frames captured at 5-minute intervals. The video recordings of the cells are shown in supplemental videos S1-S3. Each colored line represents the path of a single cell, and 30 cells were analyzed per group; (D) Quantification of cellular migration speed derived from the trajectories shown in (C); (E) Quantification of track length derived from the trajectories shown in (C). (F) Confocal microscopy images at the basal plane of STAT5A KO cells overexpressing LifeAct-mScarlet3 and ACTN1-GFP. (J) Cell size of WT, STAT5A KO, and STAT5A-rescued cells was quantified by measuring the number of pixels occupied by each cell. (H) ChIP-seq profiles of STAT5A binding at regulatory regions of ACTN1 in ARP-1 and MCF-7 cell lines. (I) Scatter plots showing the correlation between STAT5A expression and the expression of genes across different cancer cell lineages. All expression values are presented as log2(TPM+1) For Figure 4J statistical significance was calculated using one-way ANOVA followed by Dunnett’s post hoc test. Data shown are representative of at least three independent experiments.

Given the crucial role of actin organization in cell motility^22,23^, we assessed cell migration using live-cell imaging. Compared to WT cells, STAT5A KO cells exhibited slower, irregular migration patterns, characterized by shorter migration paths, reduced migration speeds, and decreased directional persistence (Figures 4C-4E; Videos S1–S3). Notably, re-expression of STAT5A not only rescued but enhanced motility beyond WT levels. This is likely due to higher-than-physiological expression of STAT5A in these cells (Fig. 2B).

To identify the role of ACTN1 in actin network disruption observed in STAT5A-deficient cells, we overexpressed ACTN1-GFP in STAT5A KO cells. Restoration of ACTN1 expression normalized cellular morphology and reestablished organized actin filament structures (Figure 4F and S3E), indicating that ACTN1 functions as a critical downstream effector mediating STAT5A-dependent maintenance of actin cytoskeletal integrity. To determine whether STAT5A directly regulates ACTN1 transcription, we analyzed available ChIP-seq datasets from the ChIP-Atlas repository. Independent studies conducted in ARP1 and MCF7 cells revealed significant STAT5A binding peaks upstream of the ACTN1 locus (Figure 4H), supporting direct transcriptional regulation. These findings suggest that the actin network defects associated with STAT5A deficiency observed in HeLa cells are conserved across different cell types. Further analysis of the DepMap RNA-seq database^24,25^, encompassing 1,518 human cancer cell lines, revealed distinct expression patterns of STAT5 paralogues. STAT5A exhibited substantial tissue-specific variability and lower median abundance compared to the more uniformly expressed STAT5B (Figure S3B). GAPDH, a well-known housekeeping gene, did not show correlation with STAT5A. in contrast, a robust correlation between STAT5A and ACTN1 expression was observed specifically in cervix- and bone-derived cell lines, consistent with our findings in HeLa cells (cervix cell lineage), while no correlation was evident in lung or myeloid cell lineages (Figure 4I).

Collectively, these data demonstrate that STAT5A deficiency disrupts the actin cytoskeleton network by downregulating critical actin-regulatory genes, and specifically ACTN1. Importantly, ACTN1 overexpression was sufficient to rescue actin organization. ChIP-seq data indicating strong STAT5A binding at the ACTN1 promoter further supports its direct role as a transcriptional regulator.

### The disrupted actin cytoskeleton in STAT5A KO cells impacts mitochondrial organization

Mitochondrial positioning depends on two complementary cytoskeletal systems: microtubules, which drive long-range transport, and actin filaments, which mediate short-range motility and local anchoring^26–28^. In STAT5A cells, we observed strong mitochondrial clustering near the nucleus (Figure 5A and 6I). To determine whether microtubule defects underlie this clustering in STAT5A KO cells, we examined the microtubule lattice by confocal microscopy. Although the network in STAT5A KO cells appeared slightly sparser—most likely because the cells are larger—it remained continuous and evenly distributed (Figure S3C) ruling out gross microtubule disorganization as the source of the phenotype. In contrast, the actin cytoskeleton network showed pronounced defects (Figure 4A). Confocal microscopy shows that in STAT5A KO HeLa cells mitochondria collapse into a dense perinuclear cluster, whereas in WT and STAT5A-rescued cells they remain evenly dispersed (Figure 5A and 6I). This mislocalisation coincides with a pronounced disassembly of the actin lattice in the KO cells (Figure 4A and 5A). In WT cells, mitochondria align along robust actin filaments and appear as short tubular structures distributed throughout the cytoplasm. By contrast, STAT5A-deficient cells contain fewer, thinner filaments, and mitochondria aggregate into compact bundles around the nucleus. Re-expression of STAT5A re-establishes both the filamentous actin network and the normal, dispersed mitochondrial morphology (Figure 5A), underscoring the role of intact actin networks as regulated by STAT5A abundance in maintaining actin integrity and, consequently, proper mitochondrial distribution. Quantification of mitochondrial-free areas revealed a significant increase in STAT5A KO cells, further highlighting the altered mitochondrial dispersal pattern (Figure 5B and 6I). Because ACTN1 emerged as the key effector linking STAT5A loss to actin disorganisation, we next asked whether restoring ACTN1 alone could also normalize mitochondrial positioning. Confocal imaging revealed that ectopic ACTN1-GFP expression in STAT5A KO cells dispersed mitochondria throughout the cytoplasm and reinstated their normal tubular morphology, mirroring the pattern seen in wild-type cells (Figure 5C). This rescue confirms that the perinuclear clustering and shape defects of mitochondria in STAT5A KO cells are secondary to the disrupted actin network and can be reversed by ectopic ACTN1 expression.

**Figure 5.**
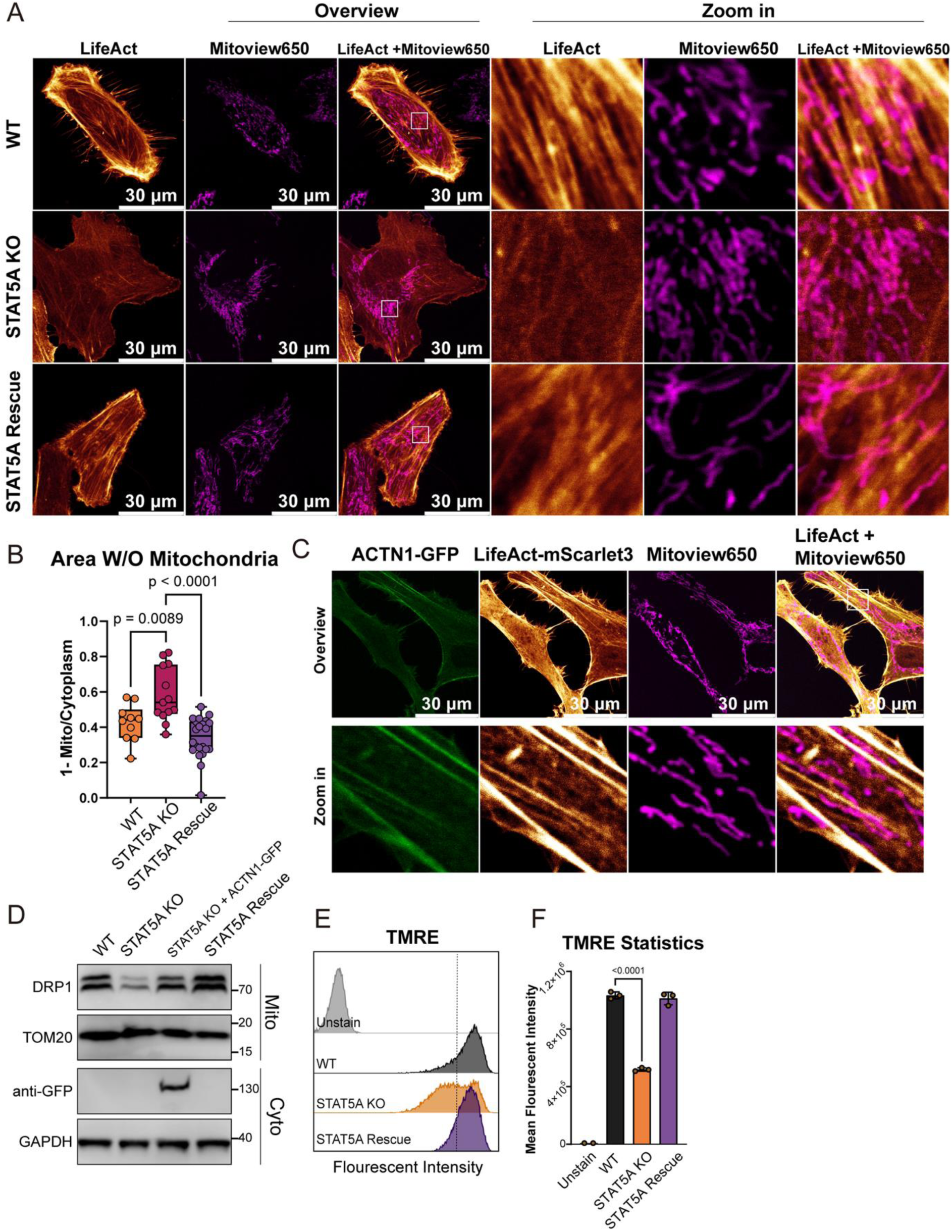
The disrupted actin cytoskeleton in STAT5A KO cells impacts mitochondrial organization. (A) Confocal microscopy images at the basal plane of WT, STAT5A KO, and STAT5A-rescued cells overexpressing LifeAct-mScarlet3 and treated with 20 nM MitoView 650. (B) Boxplot showing the fraction of mitochondrial-free cytoplasmic area as calculated by 1 minus the ratio of mitochondrial occupied to total cytoplasm area within the cells. (C) Confocal microscopy images at the basal plane of STAT5A KO cells overexpressing LifeAct-mScarlet3 and ACTN1-GFP, treated with 20 nM MitoView 650. (D) Western blot analysis of DRP1 within cytoplasmic and mitochondrial fractions taken from WT, STAT5A KO, ACTN1 overexpressing and STAT5A-rescued cells. GAPDH and TOM20 are cytoplasmic and mitochondria markers respectively. (E-F) Analysis of mitochondrial membrane potential using TMRE staining. (E) WT and STAT5A KO cells were treated with 100 nM TMRE for 20 minutes, and fluorescence intensity was quantified by flow cytometry. (F) Bar graph shows mean TMRE fluorescence intensity. For Figure 5B and 5F statistical significance was calculated using one-way ANOVA followed by Dunnett’s post hoc test. Data shown are representative of at least three independent experiments.

**Figure 6.**
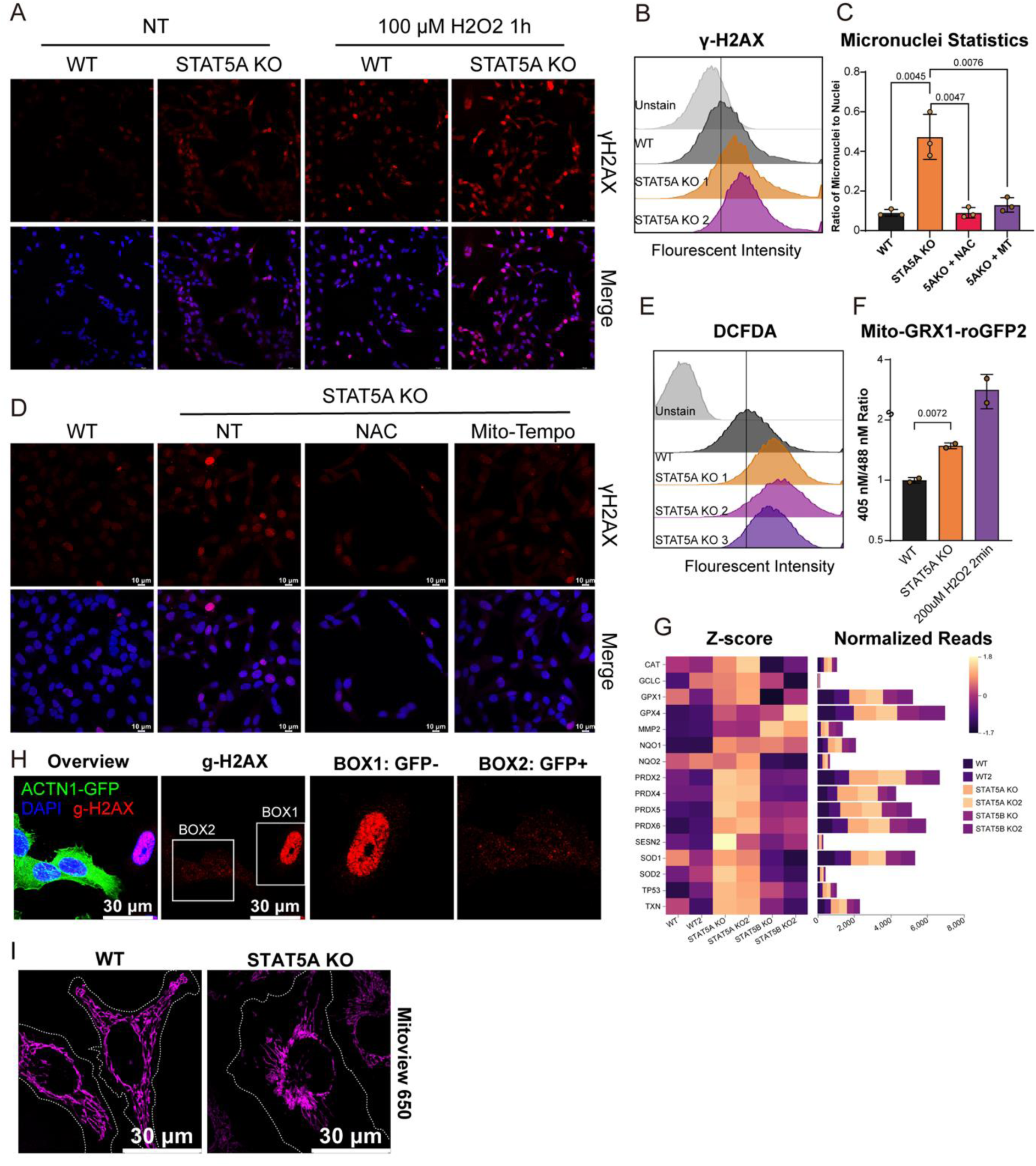
STAT5A Knockout Results in Genomic DNA Damage. (A) WT and STAT5A KO cells were either untreated or treated with 100 µM H_2_O_2_ for 1 hour. Cells were stained with anti-γH2AX (red) to detect DNA damage. (B) Quantification of γH2AX signal intensity by flow cytometry. (C) Ratio of micronuclei to nuclei in WT and STAT5A KO cells, either untreated or treated with 20 mM NAC or 100 μM Mito-TEMPO, visualized by DAPI staining. (D) WT and STAT5A KO cells, either untreated or treated with 20 mM NAC or 100 μM Mito-TEMPO, stained with anti-γH2AX (red) and DAPI (blue). (E) WT and STAT5A KO cells were treated with 10 µM DCFDA for 30 minutes, and fluorescence intensity was quantified using flow cytometry. (F) Measurement of mitochondrial ROS using mito-GRX1-roGFP2. WT and STAT5A KO cells were transfected with mito-GRX1-roGFP2 reporters, and fluorescence was analyzed by flow cytometry. The ratio of fluorescence emission at 405 nm to 488 nm excitation is presented. (G) Gene expression of ROS-inducible genes from RNA-seq analysis. Heatmap displays differential expression (z-scores) of oxidative stress-related and ROS metabolism genes in WT and STAT5A KO cells. Bar plots represent normalized read counts for these genes. (H) STAT5A KO cells overexpressing ACTN1-GFP, stained with anti-γH2AX (red) and DAPI (blue) to detect DNA damage. BOX1 shows a cell not expressing ACTN1-GFP, while BOX2 shows a cell expressing ACTN1-GFP. (I) Mitochondrial morphology of WT and STAT5A KO cells treated with MitoView 650 and Mito-TEMPO, visualized by confocal microscopy. For Figure 6C and 6F statistical significance was calculated using one-way ANOVA followed by Dunnett’s post hoc test. Data shown are representative of at least three independent experiments.

Mitochondrial clustering is a hallmark of impaired fission^29^. Because actin filaments drive fission by recruiting the dynamin-related GTPase DRP1 to the outer mitochondrial membrane^30^, we asked whether STAT5A loss hinders this recruitment. We therefore isolated mitochondria from wild-type, STAT5A KO, ACTN1 overexpression, and STAT5A-rescued cells and quantified membrane-bound DRP1 by immunoblotting (Figure 5D). DRP1 association was markedly reduced in STAT5A KO cells but restored in the ACTN1 overexpression or STAT5A rescued cells, indicating that STAT5A deficiency suppresses DRP1 recruitment depends on the ACTN1 expression. Reduced DRP1 loading is associated with mitochondrial dysfunction, which leads to mitochondrial damage and increased ROS production, ultimately impacting cellular bioenergetics^31^. We then evaluated the mitochondrial membrane potential (Δψm) using the TMRE assay, which showed a depolarization in Δψm in STAT5A KO cells compared to WT cells (Figure 5E and 5F).

Collectively, these results establish a mechanistic cascade in which STAT5A sustains ACTN1 abundance, which is crucial in maintaining actin network architecture to support cytoplasmic dispersion of mitochondria. Loss of STAT5A disassembling actin networks, driving perinuclear mitochondrial aggregation and functional impairment.

### STAT5A Knockout Results in Genomic DNA Damage

STAT5A KO cells displayed increased genomic DNA damage compared to WT cells, as demonstrated by elevated γH2AX immunofluorescence staining, a well-established nuclear marker of DNA double-strand breaks (Figure 6A)^32^. These observations were supported by flow cytometry analyses showing higher γH2AX levels and an increased frequency of micronuclei, confirming greater genomic instability in STAT5A-deficient cells (Figures 6B, 6C, and 7A). To investigate the cause of DNA damage, we focused on endogenous oxidants, which contribute to genomic instability and DNA damage responses^33^. Treatment of STAT5A KO cells with N-acetylcysteine (NAC), a broad-spectrum antioxidant, and Mito-TEMPO, a mitochondria-specific antioxidant, led to reduction in γH2AX levels (Figure 6D). Additionally, the proportion of cytosolic micronuclei, IFN and ISG gene expression in STAT5A KO cells were reduced to WT levels following Mito-TEMPO treatment (Figure S2B). These data suggest that oxidative stress related to mitochondria is the cause of DNA damage and downstream inflammatory signaling in the absence of STAT5A.

**Figure 7.**
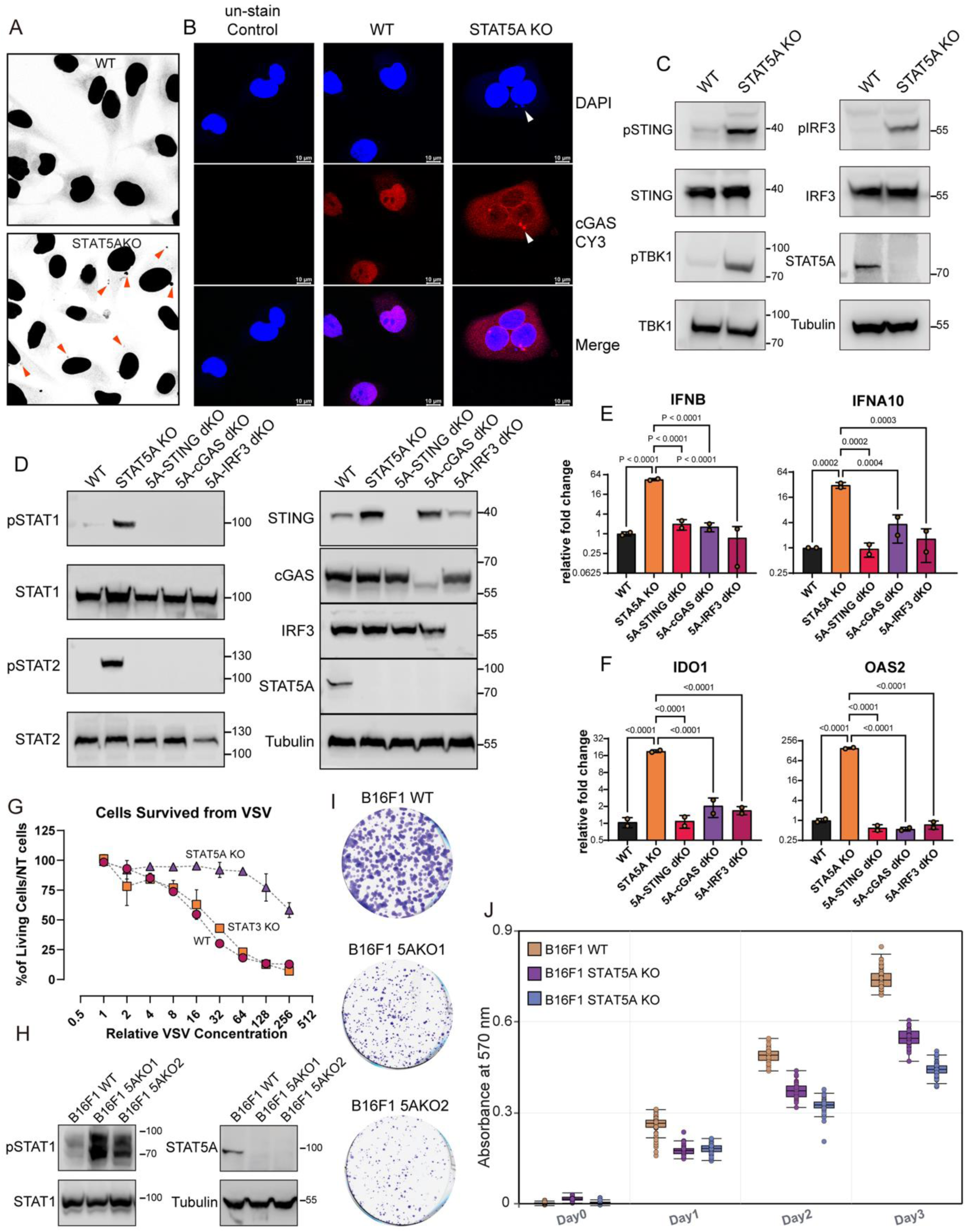
The cGAS-STING pathway is activated by cytosolic DNA in STAT5A KO cells. (A) Confocal microscopy showing DAPI staining (10 µg/mL) for micronuclei formation in WT and STAT5A KO cells. Red arrows indicate the presence of micronuclei. (B) Immunofluorescence staining showing the colocalization of cGAS with the micronuclei. DAPI stains nuclei (blue), and CY3 labels cGAS (red). (C-D) Western blot analysis of cGAS-STING pathway activation. (C) Phosphorylated STING, TBK1, and IRF3 in WT and STAT5A KO cells. (D) pSTAT1 and pSTAT2 levels in STAT5A KO, STAT5A-cGAS dKO, STAT5A-STING dKO, or STAT5A-IRF3 dKO cells. (E-F) qPCR analysis of type I IFN and ISG expression. (E) Relative expression of IFN-β and IFN-α10 in WT, STAT5A KO and STAT5A KO together with cGAS, STING, or IRF3 KO. (F) Expression levels of ISGs in WT, STAT5A KO, and STAT5A KO cells together with cGAS, STING, or IRF3 KO. (G) Percent survival after VSV exposure of WT, STAT3 KO and STAT5A KO HeLa cells relative to untreated controls after exposure to increasing concentrations of VSV for 10 hours. (H) Western blot analysis of pSTAT1 level in WT and STAT5A KO in B16F1 melanoma cells. (I) Plaque formation assay of WT and STAT5A KO B16F1 cells. (J) MTT assay measuring cell proliferation of WT and STAT5A KO B16F1 cells over three days. For Figure 7E and 7F statistical significance was calculated using one-way ANOVA followed by Dunnett’s post hoc test. Data shown are representative of at least three independent experiments

Given the observed mitochondrial dysfunction in STAT5A KO cells (Figure 5), which is frequently associated with ROS overproduction^34^, we next measured ROS levels. Using the general ROS probe DCFDA, we detected a substantial increase in fluorescence in STAT5A KO cells relative to WT, indicating elevated overall ROS levels (Figure 6E). To assess mitochondrial-specific ROS, we employed the ratiometric sensor mito-GRX1-roGFP2, which measures oxidation state by comparing fluorescence intensity at 405 nm and 488 nm^35,36^. STAT5A KO cells showed a significant increase in the 405/488 nm ratio, indicating elevated mitochondrial ROS (Figure 6F). These findings are consistent with our RNA-seq data, which revealed selective upregulation of genes involved in redox regulation and ROS metabolism—such as *CAT*, *GPX1*, *GPX4*, and *PRDX2*—in STAT5A KO but not STAT5B KO cells (Figure 6G).

Because ACTN1 was identified as a critical mediator of actin cytoskeleton and mitochondrial organization downstream of STAT5A, we examined whether restoring ACTN1 could rescue DNA damage phenotypes. Ectopic expression of ACTN1 in STAT5A KO cells significantly reduced γH2AX staining, indicating a reversal of DNA damage (Figure 6H, compare BOX2 cell with ACTN1-GFP to BOX1 cell without expression of ACTN1). To determine whether mitochondrial clustering was a consequence of elevated ROS, we treated STAT5A KO cells with Mito-TEMPO and assessed mitochondrial morphology. Despite ROS scavenging, mitochondria remained perinuclear clustered, indicating that the aberrant mitochondrial distribution arises independently of ROS levels (Figure 6I).

Collectively, these findings demonstrate that loss of STAT5A results in mitochondrial oxidative stress and increased genomic instability. The rescue of DNA damage by ACTN1 overexpression further supports the critical role of STAT5A–ACTN1 signaling in maintaining mitochondrial architecture and genomic integrity.

### The cGAS-STING pathway is activated by cytosolic DNA in STAT5A KO cells

Network correlation analysis (Figure 3C) predicted activation of cGAS and IRFs, which are key components of the cytosolic DNA sensing pathway, in STAT5A KO cells. Given the observed genomic instability in STAT5A KO cells, we investigated whether activation of the cGAS–STING pathway in this context underlies the elevated ISG expression. Using confocal microscopy with DAPI staining, we observed an increased number of micronuclei in the cytoplasm of STAT5A KO cells (Figure 7A and 6C). Subsequent immunostaining with an anti-cGAS antibody revealed that cGAS colocalized with these cytoplasmic DNA structures, suggesting activation of the cGAS-STING pathway. In WT HeLa cells, we observed that cGAS localized predominantly in the nucleus and showed minimal cytoplasmic presence (Figure 7B). This nuclear-dominant pattern of cGAS distribution is notably different from that reported in many other cell types, where cGAS often resides and functions within the cytoplasm. Such a unique localization in HeLa cells is consistent with previous studies^37^. In contrast, STAT5A KO cells displayed a marked increase in cytoplasmic cGAS (Figure 7B). The activation of the cGAS-STING pathway was further confirmed by Western blot analysis to assess the phosphorylation status of key pathway components. Elevated levels of phosphorylated STING, TBK1, and IRF3 were detected in STAT5A KO cells compared to WT controls, providing compelling evidence for pathway activation (Figure 7C).

To validate these findings, we generated knockouts of STING, cGAS, and IRF3 in STAT5A KO cells using CRISPR-Cas9. Removal any of these pathway members blocked the activation of phosphorylated STATs induced by the absence of STAT5A (Figure 7D). Concurrently, qPCR results indicated that IFN and ISG expression were restored to basal levels comparable to WT controls in double KO cells (STAT5A-STING, STAT5A-cGAS, and STAT5A-IRF3) (Figure 7E and 7F). In summary, these results demonstrate that loss of STAT5A leads to cytosolic DNA accumulation, which activates the cGAS–STING pathway.

### STAT5A KO results in an enhanced anti-viral and anti-tumor effect

Activation of the cGAS–STING pathway in tumor cells is known to induce an antiviral state and is increasingly recognized as a mechanism of anti-cancer immunity^38,39^. To assess the functional consequence of cGAS–STING activation in STAT5A KO cells, we performed a VSV infection assay in STAT5A KO, STAT3 KO, and WT cells without exogenous IFN treatment. STAT5A KO cells exhibited significantly greater resistance to VSV infection compared to both WT and STAT3 KO cells, indicating an enhanced basal antiviral state associated with STAT5A loss (Figure 7G). To explore the potential anti-cancer effects of STAT5A deficiency, we extended our analysis to the B16F1 melanoma model in both in vitro and in vivo settings. Consistent with our observations in HeLa cells, STAT5A KO in B16F1 cells led to the accumulation of pSTAT1 (Figure 7H). To determine whether STAT5A deficiency affected tumorigenic properties in vitro, we assessed the proliferation and survival rate of B16 cells cultured under standard conditions. Given that one of the hallmarks of cancer is the ability of malignant cells to sustain high proliferation and survival rates, this provided a direct measure of tumorigenicity. MTT and plaque formation assays revealed that STAT5A KO significantly reduced the proliferation rate of B16F1 cells (Figures 7I and 7J). Furthermore, preliminary in vivo experiments demonstrated enhanced infiltration of T cells (CD3+, CD8+) and dendritic cells (MHCII+CD11C+) into the tumor microenvironment of STAT5A KO B16F1 xenografts, as quantified by flow cytometry (Figure S4A). These findings suggest that the constitutive activation of the cGAS-STING pathway in STAT5A KO cells enhances anti-tumor immune responses by promoting immune cell recruitment and activation in the tumor microenvironment.

## Discussion

Our findings reveal a previously unrecognized role for STAT5A in maintaining cytoskeletal and mitochondrial homeostasis to prevent aberrant innate immune activation. STAT5A KO in HeLa cells led to pronounced disorganization of the actin cytoskeleton network, yielding larger, amorphous cells with impaired migratory dynamics. This loss of cytoskeletal integrity was accompanied by aberrant mitochondrial positioning and function, characterized by perinuclear clustering, impaired mitochondrial membrane potential, and elevated ROS production. The resulting DNA damage drove autocrine type I interferon production via the cGAS–STING pathway, resulting in constitutive activation of JAK–STAT signaling.

A key question here is to identify the driver of phenotypic outcomes associated with STAT5A KO. Ectopic expression of STAT5A reversed every one of the here observed defects in both HeLa and B16F1 cells (Figures 2, 3 and 4), indicating the crucial role of STAT5A in all these events. To clarify the order of downstream events, we dismantled the type I IFN pathway by knocking out JAK1 or the IFN-α/β receptors (IFNAR1/2) in STAT5A-KO cells. This abolished ISG up-regulation (Figure 2D and E) but did not restore expression of actin-related genes (Figure 3E and S3D), placing cytoskeletal disruption upstream of IFN signaling. Likewise, knockout of IRF3 or cGAS in STAT5A KO cells eliminated STAT1/2 phosphorylation and IFN-β production (Figure 7D and 7E), confirming that cytoplasmic-DNA sensing drives the IFN response. Our study clearly shows that the underlying reason for DNA fragmentation in STAT5A KO cells is mitochondrial ROS, as treatment with NAC or the mitochondrial antioxidant Mito-TEMPO eradicated micronuclei and γH2AX foci (Figure 6A). Yet Mito-TEMPO failed to disperse the perinuclear mitochondrial clusters characteristic of STAT5A-KO cells (Figures 6I), suggesting that mitochondrial dis-positioning precedes ROS production and is rooted in cytoskeletal defects. Indeed, STAT5A KO caused profound actin-network collapse and sharply reduced cell motility, whereas STAT5A overexpression enhanced migration beyond wild-type levels (Figure 4C-E and videos 1-3). Chromatin immunoprecipitation and transcriptomics revealed that STAT5A directly promotes transcription of ACTN1, which encodes the F-actin cross-linker α-actinin-1 (Figure 4H). Restoring ACTN1 in STAT5A KO cells rebuilt the actin lattice, redistributed mitochondria, and suppressed γH2AX accumulation (Figures 4F and 6H), tying mitochondrial stress to actin disorganization. Notably, cytoskeletal collapse persisted after Mito-TEMPO (Figure 7I) even though the IFN signature was extinguished (Figure S2B), further separating actin defects from the downstream IFN cascade. Together, these experiments place actin-network failure at the top of the hierarchy: STAT5A KO reduces ACTN1 abundance, destabilizing F-actin. The resulting cytoskeletal collapse traps mitochondria near the nucleus as well as blocks DRP1 loading (Figure 5D). This provokes mitochondrial dysfunction and ROS generation, which results in cytoplasmic DNA. This DNA engages cGAS–IRF3, unleashing type I IFN secretion and constitutive JAK–STAT signaling—completing the cascade.

The STAT5A (but not STAT5B)–cytoskeleton–mitochondria axis represents a previously unrecognized dimension of STAT5A’s function. STAT5A is classically known for regulating transcriptional programs in cell proliferation, differentiation, and survival; here we show it also serves as a structural guardian of the cell. Notably, prior studies have observed STAT5A outside the nucleus – for instance, mitochondrial-localized STAT5A can interact with metabolic enzymes to modulate respiration^40^. Our results provide a complementary perspective: rather than acting within mitochondria, STAT5A protects mitochondrial function indirectly by maintaining the actin cytoskeletal framework on which mitochondrial organization and fission depend. In the absence of STAT5A, the transcriptional network that upholds actin integrity is perturbed, leading to secondary mitochondrial dysfunction and DNA damage. This highlights STAT5A as a multifaceted regulator of cellular homeostasis, linking nuclear signaling pathways to the physical maintenance of organelle dynamics and genome stability.

Consistent with our observations, actin dynamics are now recognized as essential for mitochondrial function and cellular homeostasis^41–44^. Mitochondria rely on an intact actin cytoskeletal network for proper trafficking and localization within the cell. Disruption of this network hampers mitochondrial dynamics, leading to mitochondrial dysfunction characterized by depolarized membrane potential and increased mitochondrial ROS production^27,31,45^. STAT5A is a transcription factor of ACTN1, and upon its KO ACTN1 abundance decreases. This causes mitochondrial distribution, which is reversed by ectopic ACTN1 expression. Rather than being evenly dispersed, mitochondria in STAT5A KO cells accumulated around the nucleus, leaving the cell periphery depleted of organelles, indicating the actin disorganization in STAT5A-KO cells releases mitochondria from normal anchoring cues, causing them to coalesce in the perinuclear region. DRP1-mediated fission requires polymerized actin to achieve initial organelle constriction and recruit the fission machinery^46^. Actin filaments and myosin II form a contractile ring-like scaffold that **u**ltimately recruits DRP1 to the mitochondrial surface, where DRP1 oligomerizes and severs the organelle^15^. Our STAT5A-KO model fits well with this paradigm: disassembly of actin stress fibers in KO cells impedes DRP1’s access to mitochondria, explaining the observed hyperfused and clustered mitochondrial network. Either ectopic expression of STAT5A or ACTN1 in STAT5A KO cells rescued DRP1 levels associated to the mitochondria (Fig. 5D). Our work extends current literature by showing that ACTN1 abundance regulated by STAT5A maintains actin network integrity, governing fundamental aspects of mitochondrial cell biology.

Our study also sheds light on an emerging theme of crosstalk between the actin cytoskeleton and innate immune signaling. Recent reviews have emphasized that actin dynamics can serve as a platform for complex signaling pathways regulating type I IFN responses in cancer^47^. Our findings extend this concept to actin disorder associated cGAS activation: when the actin framework is compromised via STAT5A KO, the resulting cellular chaos – including mitochondrial dysfunction and DNA damage – provokes a DNA-driven IFN response. In essence, a healthy cytoskeleton helps silence innate immune triggers by confining organelles and genetic material to their proper place. Once this order is disturbed, cells may interpret the situation as if a pathogen or danger is present, thus activating IFN pathways. This insight places the actin cytoskeleton as a crucial integrator of structural and immune signals. It also dovetails with the notion that many pathogens actively target the host cytoskeleton^48–51^; our results suggest that, in doing so, pathogens might be trying to prevent or hijack these cytoskeleton-mediated immune alarms. Our work provides a concrete example tying actin and IFN signaling together through the intermediary of mitochondrial stress.

It has been previously suggested that STAT5 has a role in the progression of various cancers, including hematopoietic malignancies and solid tumors such as breast and prostate cancer. High expression and aberrant activation of STAT5 are associated with poor prognosis and resistance to conventional therapies^2,52,53^. Targeting STAT5 with inhibitors has shown promise in inducing apoptosis in cancer cells^7–9^. The impaired cytoskeletal and focal adhesion organization reduces cell motility, as demonstrated by live-cell imaging, where STAT5A knockout cells exhibit decreased speed and directional movement, while higher STAT5A abundance increases them. By disrupting cell motility and invasion, STAT5A deficiency may hinder metastasis, which is critical in cancer progression^54,55^. These structural abnormalities have profound functional consequences. Additionally, the activation of the cGAS-STING pathway in STAT5A-deficient cells leads to enhanced type I IFN responses, which can promote anti-tumor immunity by activating immune cells and inducing tumor cell apoptosis^38,56,57^. Indeed, we show that B16F1 melanoma cells grow slower in vitro and in mouse xenograft, recruiting immune cells upon STAT5A KO (Figure 7 and S4A). This immune activation adds another layer to the anti-cancer effects of STAT5A deficiency, suggesting that targeting STAT5A could offer a multifaceted therapeutic strategy in cancer treatment.

Neurodegenerative diseases are another arena where actin–mitochondria communication is critical. Neurons rely on a delicately balanced transport system to deliver mitochondria to synapses and areas of high energy demand. Disruption of actin stability in neurons is known to cause mitochondrial mislocalization and energy deficits, contributing to neurodegenerative pathology^15^. In this light, it is worth considering whether STAT5A or ACTN1 dysregulation in the nervous system could play a role in diseases like ALS or Alzheimer’s, where both the cytoskeleton and mitochondrial health are compromised. For example, abnormal cofilin–actin aggregates in neurons can induce mitochondrial fragmentation and oxidative stress, leading to synaptic loss and cell death^15,58,59^. An analogous loss of actin regulatory control (potentially through impaired STAT5A signaling or actinin function) might similarly tip the balance toward mitochondrial failure and neuronal injury. Future studies in model organisms and patient samples could investigate whether variations in STAT5A–ACTN1 activity correlate with neurodegenerative disease progression or severity.

In summary, our work defines ACTN1 abundance as controlled by the STAT5A transcription factor as a pivotal regulator of cytoskeleton-dependent mitochondrial homeostasis. Through transcriptional controlling ACTN1 expression among other actin regulators, STAT5A ensures that actin filaments form a robust network that supports mitochondrial fission, distribution, and functionality. This connection between a signaling mediator (STAT5A), a structural protein (α-actinin-1), and organelle behavior (mitochondrial dynamics and ROS production) broadens the paradigm of how cellular architecture is governed. It illustrates that safeguarding genomic stability is not solely the domain of DNA repair pathways but also relies on maintaining the infrastructure (cytoskeletal and organellar) that prevents damage from arising in the first place. The correlation between STAT5A levels and ACTN1 gene expression, as found in many different organelles (but not all, Figure 4I) suggests the biological relevance of our findings, which open new questions about cytoskeletal coordination of organelle biology and its regulation by signaling pathways. Given the emerging appreciation of actin– mitochondria interactions in health and disease, the STAT5A–ACTN1 axis identified here could represent a novel therapeutic target or biomarker in conditions ranging from cancer to neurodegeneration. Ultimately, this study reinforces the concept that cellular homeostasis is a multi-layered process, requiring tight integration of signaling, transcription, cytoskeletal dynamics, and organelle function to preserve the integrity of the cell.

## Materials and Methods

The source of all materials used in this study are detailed in Table S1.

### Cell Culture

HeLa, B16F1 and HEK293T cells were cultured in Dulbecco’s modified Eagle’s medium (DMEM) supplemented with 10% fetal bovine serum (FBS), 1% Pyruvate and 1% penicillin-streptomycin. All the cells were routinely maintained in a humidified incubator at 37°C with 5% CO_2_.

### Gene Knockout

Gene knockout cell lines were generated using the CRISPR/Cas9 system as described previously^60^. STAT1, STAT2, and STAT3 knockout cell lines were prepared earlier^61^. For cGAS, STING, and IRF3 knockout, plasmids were obtained from Addgene (details provided in the materials list). For the generation of STAT4, STAT5A, STAT5B, and STAT6 knockout cell lines, target-specific oligonucleotides were designed and cloned into the PX459 vector. This vector was then transfected into wild-type cells. Forty-eight hours post-transfection, positive cells were selected either by fluorescence-activated cell sorting (FACS) or puromycin selection, followed by culturing as single-cell clones. Each monoclonal cell line was validated for successful gene knockout using immunoblotting assays.

The guide RNA (gRNA) sequences used for each gene are listed below:

STAT4: AACAGATGCCGAATTTCCAT

STAT5A: AGTAGTGCCGGACCTCGATG

STAT5B: GTTCATTGTACAATATATGG

STAT6: GCTGGGAATAAATGTCCACC

### Lentivirus Preparation and Stable Cell Line Generation

Lentivirus Production: HEK293T cells were seeded in 10 cm culture dishes to achieve 70–80% confluency at the time of transfection. The following plasmids were used for transfection: transfer vector containing the gene of interest (1.64 pmol), psPAX2 (1.3 pmol), and pMD2.G (0.72 pmol). Transfection was performed using jetPrime or jetPEI reagents, following the manufacturer’s protocols. Approximately 3 hours post-transfection, the medium was replaced with fresh culture medium. The culture supernatant, containing lentiviral particles, was collected 24–48 hours post-transfection, depending on cell growth conditions. The supernatant was then filtered through a 0.45 µm filter to remove cellular debris and stored at –80°C until use.

Stable Cell Line Generation: Target cells were seeded in 6-well plates and infected with lentiviral particles in the presence of 8 µg/mL polybrene to enhance transduction efficiency. After 24 hours, the infection medium was replaced with fresh growth medium. Forty-eight hours post-infection, cells were subjected to selection using puromycin at a concentration of 1 µg/mL for 5–7 days. Individual colonies or bulk populations of antibiotic-resistant cells were then expanded for further analysis.

### Immunoblotting Assay

For the immunoblotting assay, cells were lysed using RIPA buffer (50 mM Tris-HCl pH 7.4, 150 mM NaCl, 1% NP-40, 0.5% sodium deoxycholate, 0.1% SDS) supplemented with protease and phosphatase cocktail inhibitors. The lysates were incubated on ice for 5 minutes, followed by centrifugation at 14,000 rpm for 15 minutes at 4°C to remove insoluble debris. Protein concentrations were determined using the BCA assay. Equal amounts of protein were mixed with 4× SDS sample buffer and boiled at 95°C for 5 minutes. Proteins were separated by SDS-PAGE on polyacrylamide gel and then transferred to NC membranes. Membranes were blocked with 5% BSA in TBST (20 mM Tris-HCl pH 7.5, 150 mM NaCl, 0.1% Tween-20) for 1 hour at room temperature, followed by overnight incubation at 4°C with primary antibodies diluted in the same buffer. After washing, membranes were incubated with HRP-conjugated secondary antibodies for 1 hour at room temperature. Protein bands were visualized using enhanced chemiluminescence (ECL) reagents and imaged with a chemiluminescent imaging system.

### RNA isolation and quantitative reverse transcription PCR

The extraction of total RNA from cells was performed using TRIzol reagent according to the manuscript. For mRNA detection, cDNA was synthesized from 1 μg of total RNA with the qScript cDNA Synthesis Kit. Quantitative polymerase chain reaction (qPCR) was performed using ChamQ SYBR qPCR Master Mix. Quantitation of the relative expression levels of target genes was performed in triplicate and calculated using the 2-ΔΔCT method. GAPDH was used as endogenous controls for normalization. PCR primers of indicated genes are shown in Table S2.

### Bulk RNA-seq

After the indicated treatments, cells were collected for total RNA isolation using the NucleoSpin RNA II Mini Spin Kit. Library preparation and sequencing were performed by The Mantoux Bioinformatics Institute at the Nancy and Stephen Grand Israel National Center for Personalized Medicine, Weizmann Institute of Science. Initial data analysis, including quality control, mapping, and read counting, was carried out using the UTAP platform^62^. Specifically, Reads were trimmed using cutadapt^63^ (parameters: -a AGATCGGAAGAGCACACGTCTGAACTCCAGTCAC -a “A{10}” –times 2 -u 3 -u -3 -q 20 -m 25). Reads were mapped to genome hg38 using STAR^64^ v2.5.2b (parameters: –alignEndsType EndToEnd, –outFilterMismatchNoverLmax 0.05, –twopassMode Basic, –alignSoftClipAtReferenceEnds No). The pipeline quantifies the 3’ of RefSeq annotated genes (The 3’ region contains 1,000 bases upstream of the 3’ end and 100 bases downstream): We used the 3’ end (1000bp) of the transcripts for counting the number of reads per gene. Counting (UMI counts) was done after marking duplicates (in-house script) using HTSeq-count^65^ in union mode. Normalization of the counts and differential expression analysis was performed using DESeq2^66^ with the parameters: betaPrior=True, cooksCutoff=FALSE, independentFiltering=FALSE. Raw P values were adjusted for multiple testing using the procedure of Benjamini and Hochberg^67^. Subsequent downstream analyses were performed using the QIAGEN Ingenuity Pathway Analysis platform or custom Python scripts.

### Immunofluorescence

For immunofluorescence (IF), cells grown on glass-bottom chambers were treated as specified. For cGAS and γH2AX staining, cells were fixed with 2% paraformaldehyde for 15 minutes at room temperature, followed by permeabilization with 0.1% Triton X-100 for 15 minutes. For cytoskeleton staining, cells were fixed with 3% paraformaldehyde in PBS containing 0.25% Triton X-100 and 0.2% glutaraldehyde for 15 minutes at room temperature. After fixation, cells were treated sequentially with 10 mg/mL sodium borohydride in PBS for 10 minutes and 2 M glycine in PBS for an additional 10 minutes to quench autofluorescence. Subsequently, cells were incubated with the specified primary antibodies for 30 minutes at room temperature, followed by staining with fluorophore-conjugated secondary antibodies for immunofluorescence detection. Nuclei were counterstained with 1 µg/mL DAPI solution. Samples were imaged using a DMi8 Leica (Leica Microsystems, Mannheim, Germany) confocal laser-scanning microscope, equipped with a pulsed white-light and 405 nm lasers using a HC PL APO 63x/1.2 water-immersion objective and HyD SP GaAsP detectors. For micronuclei detection, cells were fixed and stained with DAPI at a concentration of 10 µg/mL. Image intensities were normalized by dividing each pixel’s intensity by the 95th percentile intensity of the corresponding image and were subsequently displayed as a reversed grayscale.

### Live Cell Imaging

Live cell imaging was conducted to monitor cellular dynamics over time. GFP-stably labeled cells (3,000 cells per well) were seeded in imaging chambers (IBIDI GMBH, 80841) and allowed to adhere overnight under standard culture conditions (37°C, 5% CO₂). Imaging was performed in a temperature- and CO₂-controlled environment (Okolab, Pozzuoli, Italy) using a Nikon Eclipse TI-E inverted microscope (Nikon Corp.) equipped with a CFI Super Plan Fluor ELWD ADM 40×/0.6 objective, a GFP fluorescence filter cube, and a cooled electron-multiplying charge-coupled device (EMCCD) camera (iXon Ultra 888; Andor). Cell tracking and speed analysis were performed using spot detection algorithms in Imaris software (Bitplane, Oxford Instruments).

### Flow Cytometry for pSTAT and γ-H2AX Detection

HeLa cells were harvested after treatment and fixed with 2% paraformaldehyde (PFA) for 15 minutes at room temperature to preserve phosphorylation states. Residual PFA was quenched by treating the cells with 2M glycine for 20 minutes at room temperature. Following fixation, the cells were permeabilized with methanol for 30 minutes at room temperature. After permeabilization, the cells were washed twice with 1% BSA/PBS to remove any remaining methanol. For staining, the cells were incubated with fluorochrome-conjugated antibodies for 30 minutes at room temperature in the dark. After incubation, the cells were washed twice with 1% BSA/PBS to remove any unbound antibodies. Finally, the cells were resuspended in 1% BSA/PBS, analyzed using CytoFlex systems (Beckman Coulter). Datasets were analyzed using CytExpert and FlowJo V10.

### Flow Cytometry for Mito-GRX1-roGFP2 Detection

To detect Mito-GRX1-roGFP2 expression via flow cytometry^36,68,69^, HeLa cells were first transfected with the pLPCX Mito-GRX1-roGFP2 plasmid at a concentration of 2 µg per well in a 6-well plate. 16 hours after transfection, the cells were harvested and resuspended in non-phenol red DMEM containing 2% FBS. The cells were then immediately subjected to flow cytometric analysis to maintain the redox-sensitive properties of the Mito-GRX1-roGFP2 sensor. During analysis, laser excitation at 405 nm and 488 nm wavelengths was used to detect the oxidized and reduced states of the roGFP2 probe, respectively. The ratio of fluorescence intensities was calculated to assess the mitochondrial redox state in the cell population. Samples were analyzed using CytoFlex systems (Beckman Coulter). Datasets were analyzed using CytExpert and FlowJo V10.

### Mitochondria stain and Dispersal Ratio analysis

For mitochondrial staining, cells were incubated with MitoView650 or Mitotracker Green FM at a final concentration of 20 nM, and nuclei were stained with Hoechst 33342 at a final concentration of 1 µg/mL. Both stains were applied for 30 minutes at 37°C in a humidified incubator with 5% CO2. Following incubation, the cells were thoroughly washed with pre-warmed non-phenol red DMEM to remove any excess dye. Images of the cells were captured using a confocal microscope, utilizing bright field, DAPI, and Mito-Green channels. The mitochondrial dispersal ratio was then analyzed using a custom Python program, which detected pixel areas corresponding to the whole cell, mitochondria (green), and nuclei (blue) regions. The mitochondrial dispersal ratio was calculated using the formula: (green area pixels) / (whole cell pixels - blue area pixels).

### VSV infection and Crystal Violet Stain

For VSV infection, HeLa WT and mutation cells were first seeded into a 96-well plate at a density of 20,000 cells per well and allowed to adhere overnight. The following day, after the indicated treatments, the cells were infected with VSV and incubated for the specified time, depending on the experimental requirements. To assess cell viability and visualize virus-induced cytopathic effects, the cells were fixed with 2% PFA, and residual PFA was quenched with 2M glycine. Cells were then stained with crystal violet solution for 5 minutes. Excess stains were removed by washing the wells with water, and the plates were left to air-dry. Absorbance was measured at 590 nm using a plate reader, and the survival ratio was calculated by comparing the OD of the treated group to the non-infected control group.

### Anti-proliferation Assay

Anti-proliferation assays were performed as previously described^70^. Briefly, HeLa cells (2000 per well) were seeded into flat-bottom, 96-well plates and allowed to adhere overnight. The following day, cells were treated with a series of threefold serial dilutions of IFN-β, starting from 10 nM. After 96 hours of incubation, cell viability was evaluated by crystal violet staining, same as described before.

### Tumor Growth In Vivo and Flow Cytometry analysis

B16 cells (4 × 10^5 cells/100 μL in media) were injected orthotopically into a Left Flank of C57BL/6 female mice. Mice were sacrificed on day 10 and subsequently local tumors were removed for further analysis. Primary tumors cut into small pieces and crushed with a syringe plunger were passed through a 40 μm filter to remove any remaining large particles from the single cell suspension. The cells were stained with LIVE DEAD Staining (Aqua Zombie) 1;1000 dilution 100ul per sample for 30 mins on ice and washed with FACS buffer (1% BSA, 0.05mM EDTA, PBS) for 10 mins. Cells were blocked with the CD16/CD32 monoclonal antibody and stained with specific antibodies for 1 hr. Samples were analyzed using CytoFlex systems (Beckman Coulter). Datasets were analyzed using CytExpert and FlowJo V10.

### Statistical analysis

All experiments were independently performed at least three times. The values are presented as the mean ± standard deviation (SD). The analysis was performed using GraphPad Prism (Prism 9 for Windows). The significance of differences between groups was evaluated using unpaired one-tailed Student’s T-tests for comparisons involving two groups. For multiple group comparisons to a specific group, one-way analysis of variance (ANOVA) was used, followed by Dunnett’s correction. The P-value of less than 0.05 was considered statistically significant.

## RESOURCE AVAILABILITY

### Lead contact

Further information and requests for resources and reagents should be directed to and will be fulfilled by Gideon Schreiber.

### Materials availability

Materials included in this manuscript will be shared upon request.

### Data availability

The RNA-seq datasets generated and analyzed for this study are available at GSE285030. These data are publicly available as of the date of publication.

Any additional information required to reanalyze the data reported in this paper is available from the lead contact upon request.

## ACKNOWLEDGMENTS

We would like to express our sincere gratitude to the following individuals for their invaluable support and contributions to this research: Ze Hong (Nanjing University of Chinese Medicine, Nanjing) for his expert guidance on the cGAS experiments. Dana Reichmann (Hebrew University, Jerusalem), Tslil Ast and Zvulun Elazar (Weizmann Institute of Science, Rehovot) for their mentorship in mitochondria experiments and for generously sharing essential materials. Sabina Winograd-Katz (Weizmann Institute of Science, Rehovot) for providing critical materials and for expert guidance related to cytoskeleton organization and visualization. Moshe Goldsmith (Weizmann Institute of Science, Rehovot) for his invaluable assistance on the flow cytometry instrument. Their expertise and support have been instrumental in the successful completion of this study. This work was supported by a grant from Minerva Foundation number 714144 and Israel Cancer Research Foundation number 1344142.

## AUTHOR CONTRIBUTIONS

G.S and B.S developed the study concept and research questions. G.S supervised the project. B.S designed the methods, performed experiments and gathered data throughout the manuscript. M. K performed the B16F1 and mouse experiments. R.N helped with the microscopy work. E.Z helped in preparing materials and participated in data analysis. B.G guided the cytoskeleton work and provided crucial materials. B.S, M. K, B.G and G.S analyzed and interpreted the data. B.S wrote the initial draft of the manuscript, created figures, tables and videos. B.S, M. K, R.N, E.Z, B.G and G.S reviewed and revised the manuscript.

## DECLARATION OF INTERESTS

### Declaration of generative AI and AI-assisted technologies in the writing process

During the preparation of this work the authors used ChatGPT in order to improve the readability and language of the manuscript. After using this tool/service, the authors reviewed and edited the content as needed and take full responsibility for the content of the published article.

## Supplemental Information

**Figure S1.**
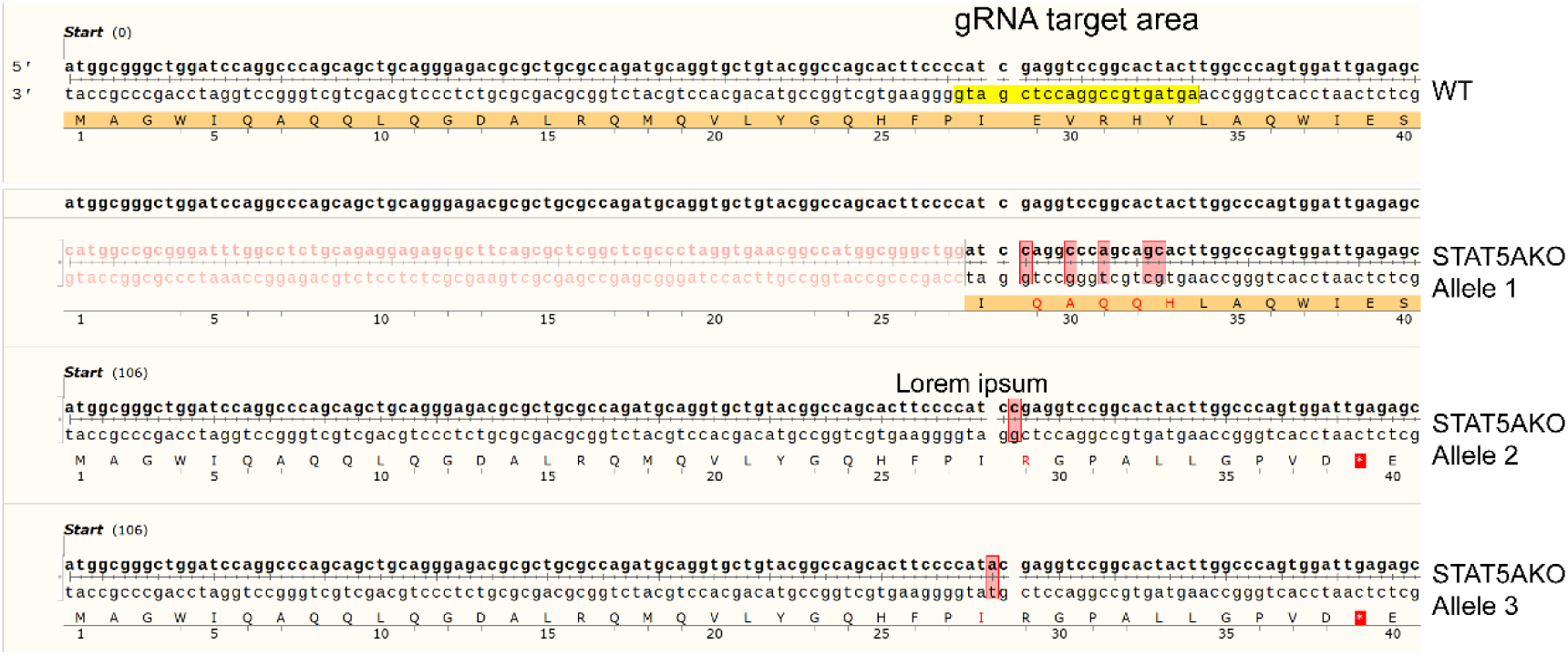
Confirmation of STAT5A Knockout in HeLa Cells by. DNA Sequencing Verification of STAT5A KO HeLa Cells. The gRNA target area is indicated in the WT sequence (top panel) and highlighted in yellow. STAT5A KO alleles (Allele 1, Allele 2, and Allele 3) exhibit CRISPR-Cas9-induced mutations in the target region. Allele 1: Large deletion including starting codon. Allele 2: 1 base pair insertion. Allele 3: 1 base pair deletion.

**Figure S2.**
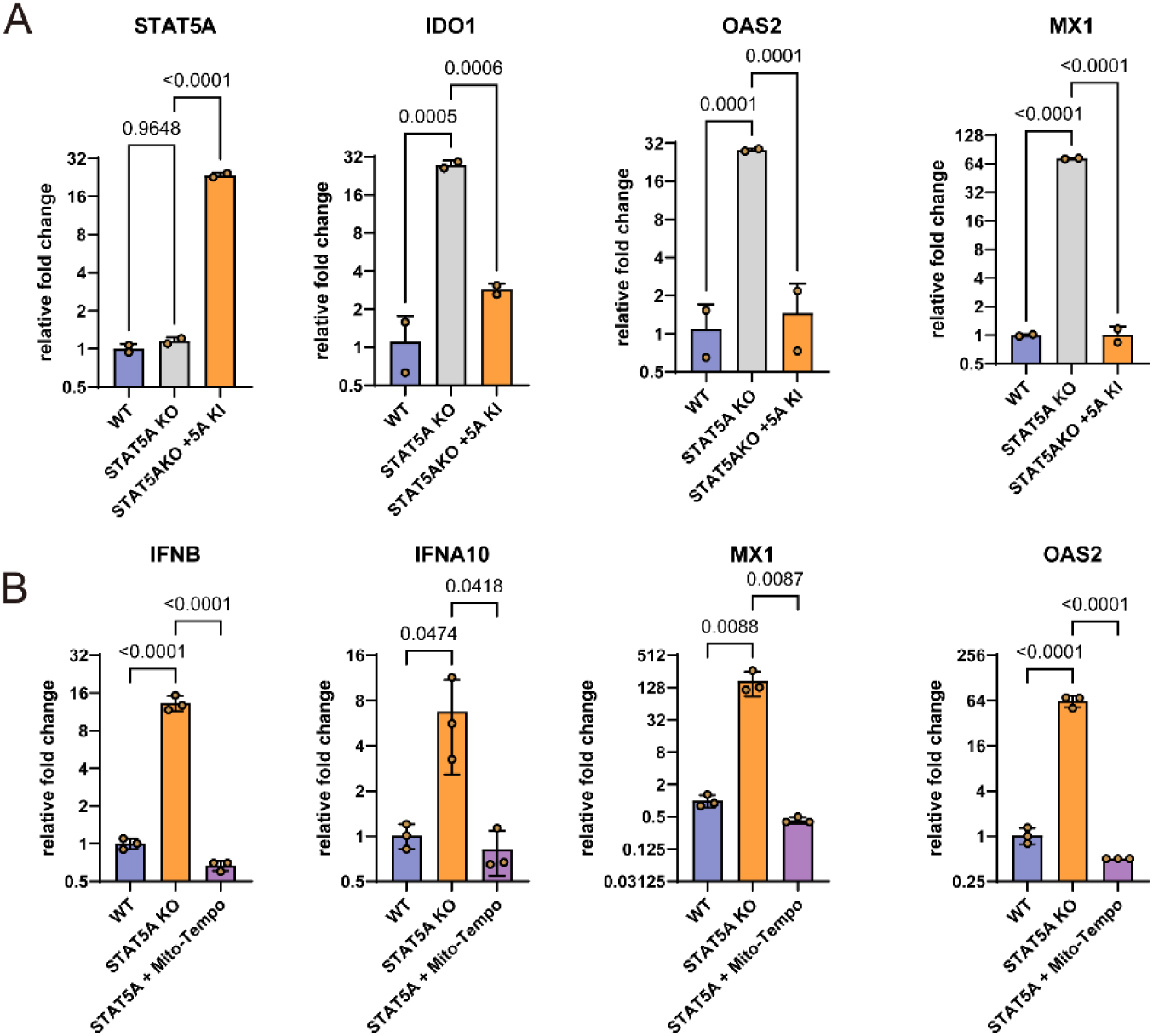
ISG and type I IFN gene expression abundance. (A) ISG expression abundance in STAT5A Rescue Cells. Relative expression abundance of STAT5A and ISGs in WT, STAT5A, and STAT5A rescue (STAT5A KO + STAT5A Knock In (KI) cells determined by quantitative qPCR. (B) IFN and ISG expression in STAT5A KO cells after two passages with 200 μM Mito-TEMPO treatment as determined by qPCR. Statistical significance was calculated using one-way ANOVA followed by Dunnett’s post hoc test. Error bars indicate standard deviation. Data shown are representative of at least three independent experiments.

**Figure S3.**
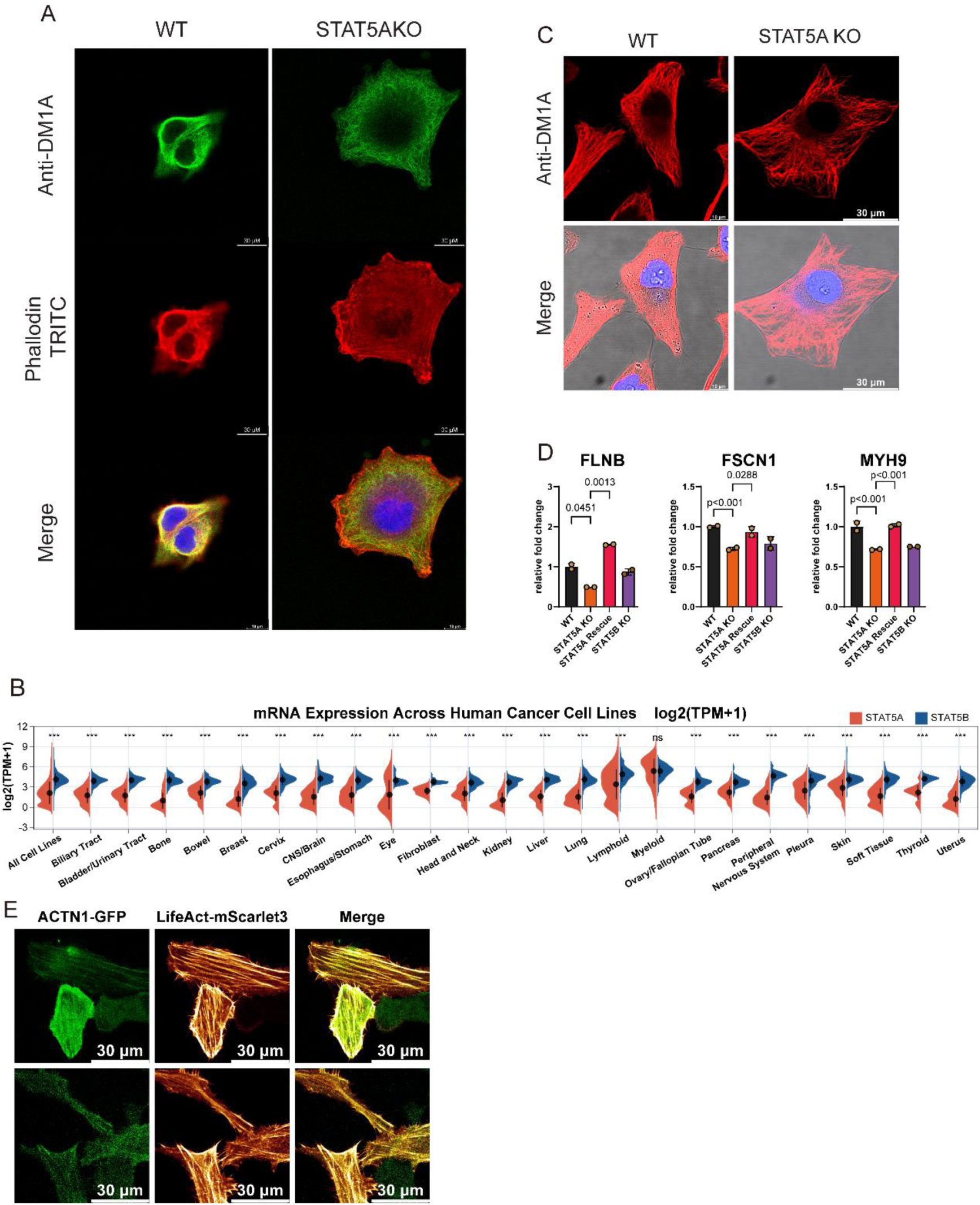
Related to Figure 4. (A) Immunofluorescence images of WT and STAT5A KO B16F1 cells showing cytoskeletal organization. Microtubules were stained with anti-DM1A (green), and F-actin was stained with TRITC-conjugated phalloidin (red). (B) Violin plots illustrating mRNA expression of STAT5A and STAT5B across various human cancer cell lines, shown as log2(TPM+1). (D) Immunofluorescence images of WT and STAT5A KO HeLa cells showing microtubules organization stained with anti-DM1A. (D) qPCR analysis of cytoskeleton and adhesion genes expression in WT, STAT5A KO, STAT5A Rescue cells. (E) Confocal microscopy images at the basal plane of STAT5A KO cells overexpressing LifeAct-mScarlet3 and ACTN1-GFP. For figure S3D, statistical significance was calculated using one-way ANOVA followed by Dunnett’s post hoc test. Error bars indicate standard deviation. Data shown are representative of at least three independent experiments.

**Figure S4.**
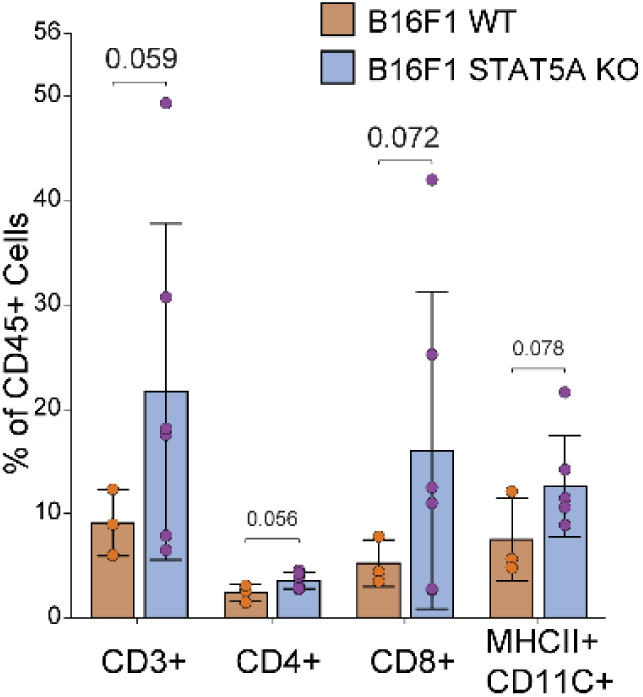
Related to Figure 7. Flow cytometric quantification of T-cell (CD3+, CD4+, CD8+) and dendritic cell (MHCII+CD11C+) infiltration in the tumor microenvironment of B16F1 xenografts mouse model as determined by FACS.

**Video S1-S3.** Live-cell imaging of cell migration associated with Figure 4C-E. Cell migration was observed using live-cell imaging of WT STAT5A and STAT5A rescue HeLa cells labeled with GFP via lentiviral transfection. Migration was visualized using an inverted microscope at 40× magnification. The time-lapse video, spanning 18 hours, is compressed into 8 seconds.

**Table s1.**
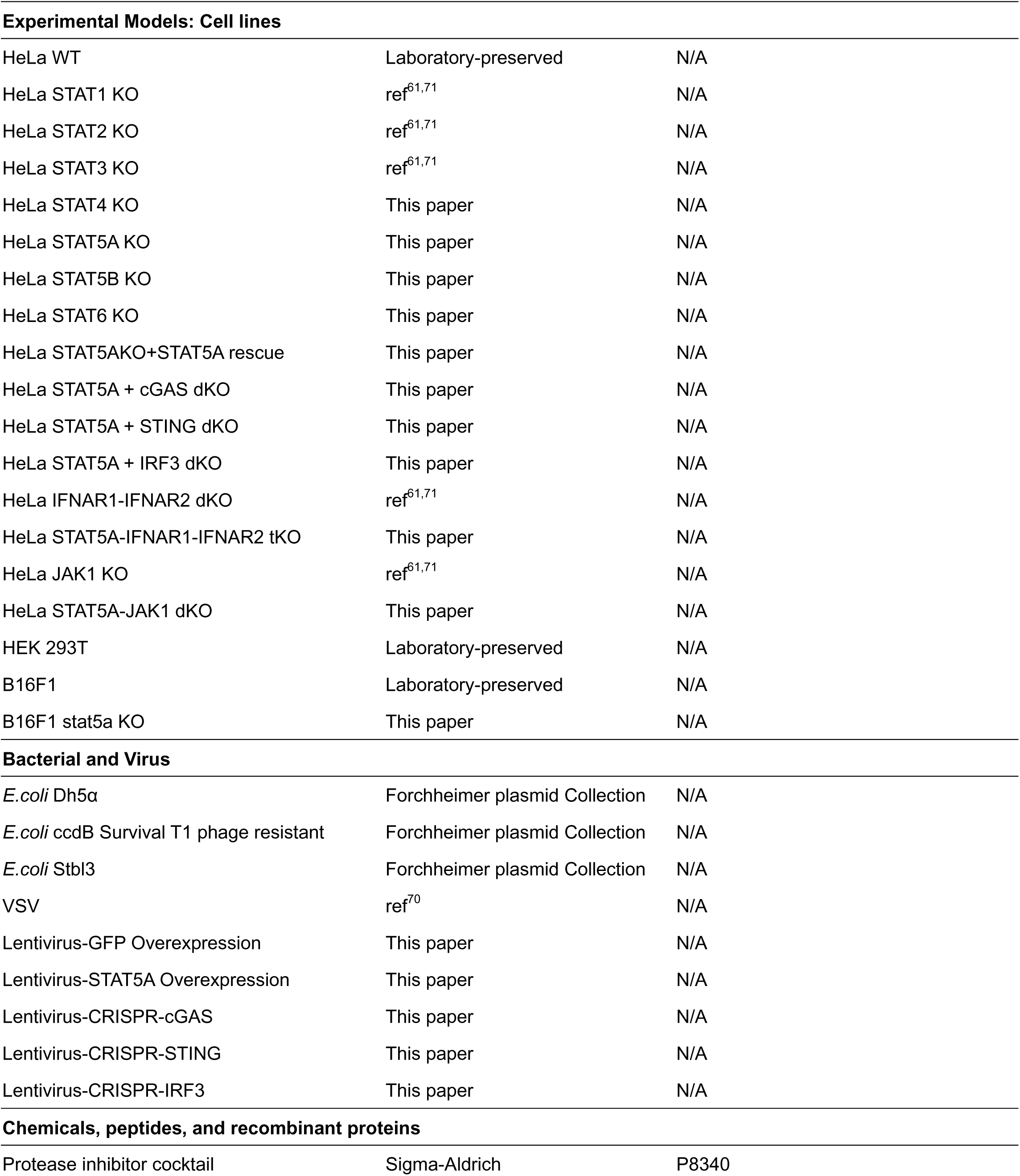
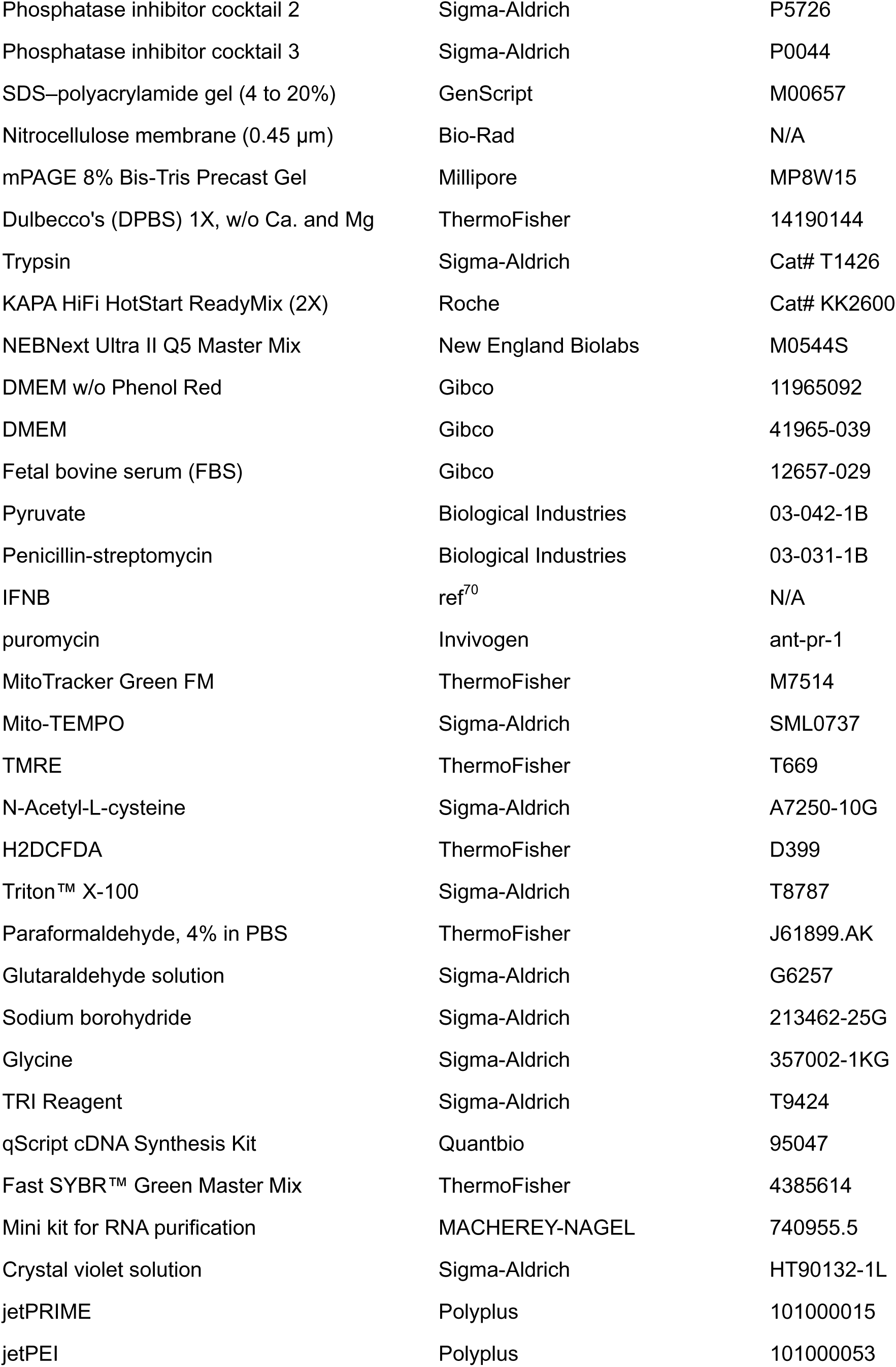
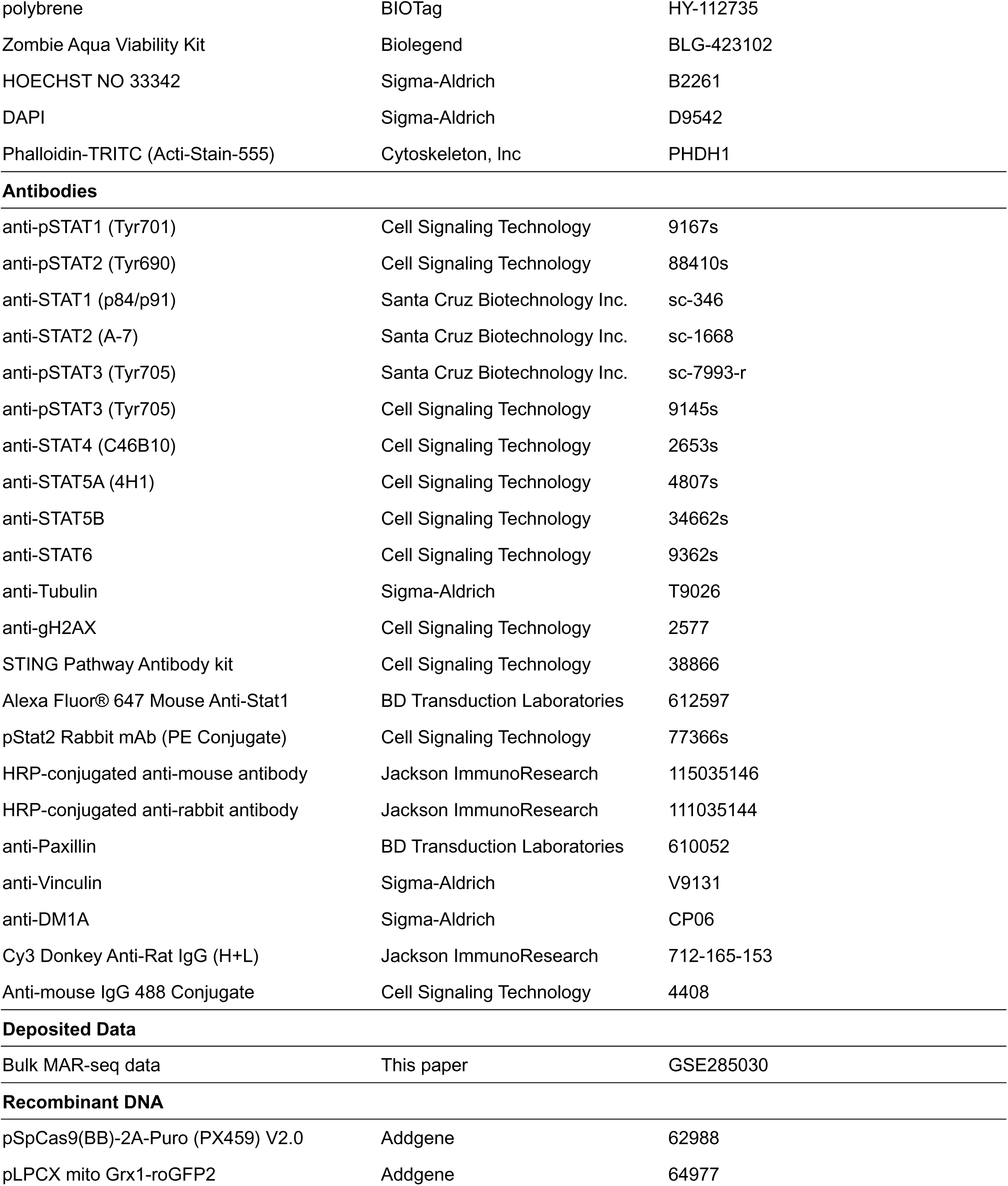
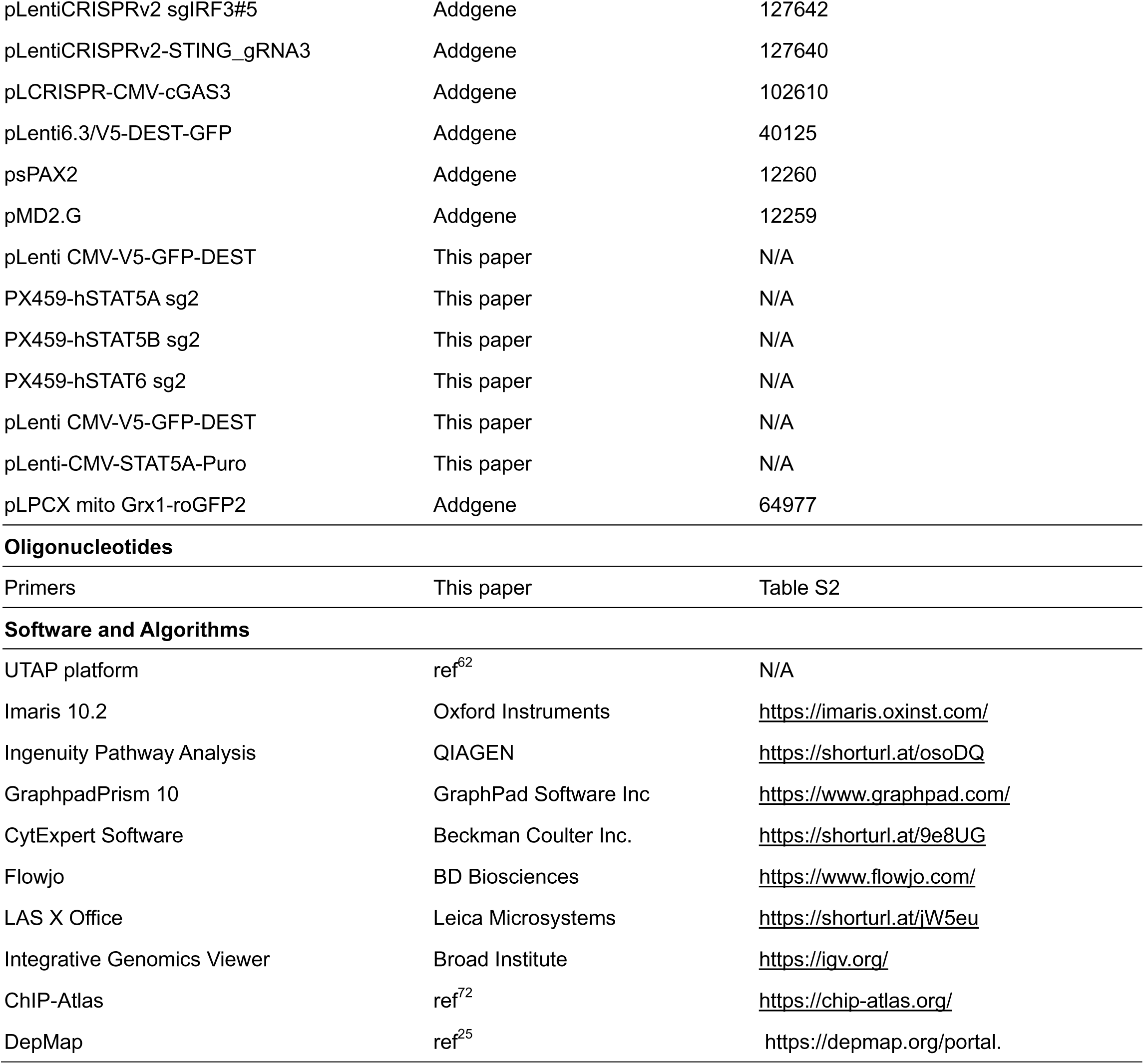
REAGENT and RESOURCE.

| Table S1. REAGENT and RESOURCE | SOURCE | IDENTIFIER |
| --- | --- | --- |
| **Experimental Models: Cell lines** |  |  |
| HeLa WT | Laboratory-preserved | N/A |
| HeLa STAT1 KO | ref<sup>61,71</sup> | N/A |
| HeLa STAT2 KO | ref<sup>61,71</sup> | N/A |
| HeLa STAT3 KO | ref<sup>61,71</sup> | N/A |
| HeLa STAT4 KO | This paper | N/A |
| HeLa STAT5A KO | This paper | N/A |
| HeLa STAT5B KO | This paper | N/A |
| HeLa STAT6 KO | This paper | N/A |
| HeLa STAT5AKO+STAT5A rescue | This paper | N/A |
| HeLa STAT5A + cGAS dKO | This paper | N/A |
| HeLa STAT5A + STING dKO | This paper | N/A |
| HeLa STAT5A + IRF3 dKO | This paper | N/A |
| HeLa IFNAR1-IFNAR2 dKO | ref<sup>61,71</sup> | N/A |
| HeLa STAT5A-IFNAR1-IFNAR2 tKO | This paper | N/A |
| HeLa JAK1 KO | ref<sup>61,71</sup> | N/A |
| HeLa STAT5A-JAK1 dKO | This paper | N/A |
| HEK 293T | Laboratory-preserved | N/A |
| B16F1 | Laboratory-preserved | N/A |
| B16F1 stat5a KO | This paper | N/A |
| **Bacterial and Virus** |  |  |
| *E.coli* Dh5α | Forchheimer plasmid Collection | N/A |
| *E.coli* ccdB Survival T1 phage resistant | Forchheimer plasmid Collection | N/A |
| *E.coli* Stbl3 | Forchheimer plasmid Collection | N/A |
| VSV | ref<sup>70</sup> | N/A |
| Lentivirus-GFP Overexpression | This paper | N/A |
| Lentivirus-STAT5A Overexpression | This paper | N/A |
| Lentivirus-CRISPR-cGAS | This paper | N/A |
| Lentivirus-CRISPR-STING | This paper | N/A |
| Lentivirus-CRISPR-IRF3 | This paper | N/A |
| **Chemicals, peptides, and recombinant proteins** |  |  |
| Protease inhibitor cocktail | Sigma-Aldrich | P8340 |
| Phosphatase inhibitor cocktail 2 | Sigma-Aldrich | P5726 |
| Phosphatase inhibitor cocktail 3 | Sigma-Aldrich | P0044 |
| SDS–polyacrylamide gel (4 to 20%) | GenScript | M00657 |
| Nitrocellulose membrane (0.45 µm) | Bio-Rad | N/A |
| mPAGE 8% Bis-Tris Precast Gel | Millipore | MP8W15 |
| Dulbecco's (DPBS) 1X, w/o Ca. and Mg | ThermoFisher | 14190144 |
| Trypsin | Sigma-Aldrich | Cat# T1426 |
| KAPA HiFi HotStart ReadyMix (2X) | Roche | Cat# KK2600 |
| NEBNext Ultra II Q5 Master Mix | New England Biolabs | M0544S |
| DMEM w/o Phenol Red | Gibco | 11965092 |
| DMEM | Gibco | 41965-039 |
| Fetal bovine serum (FBS) | Gibco | 12657-029 |
| Pyruvate | Biological Industries | 03-042-1B |
| Penicillin-streptomycin | Biological Industries | 03-031-1B |
| IFNB | ref<sup>70</sup> | N/A |
| puromycin | Invivogen | ant-pr-1 |
| MitoTracker Green FM | ThermoFisher | M7514 |
| Mito-TEMPO | Sigma-Aldrich | SML0737 |
| TMRE | ThermoFisher | T669 |
| N-Acetyl-L-cysteine | Sigma-Aldrich | A7250-10G |
| H2DCFDA | ThermoFisher | D399 |
| Triton™ X-100 | Sigma-Aldrich | T8787 |
| Paraformaldehyde, 4% in PBS | ThermoFisher | J61899.AK |
| Glutaraldehyde solution | Sigma-Aldrich | G6257 |
| Sodium borohydride | Sigma-Aldrich | 213462-25G |
| Glycine | Sigma-Aldrich | 357002-1KG |
| TRI Reagent | Sigma-Aldrich | T9424 |
| qScript cDNA Synthesis Kit | Quantbio | 95047 |
| Fast SYBR™ Green Master Mix | ThermoFisher | 4385614 |
| Mini kit for RNA purification | MACHEREY-NAGEL | 740955.5 |
| Crystal violet solution | Sigma-Aldrich | HT90132-1L |
| jetPRIME | Polyplus | 101000015 |
| jetPEI | Polyplus | 101000053 |
| polybrene | BIOTag | HY-112735 |
| Zombie Aqua Viability Kit | Biolegend | BLG-423102 |
| HOECHST NO 33342 | Sigma-Aldrich | B2261 |
| DAPI | Sigma-Aldrich | D9542 |
| Phalloidin-TRITC (Acti-Stain-555) | Cytoskeleton, Inc | PHDH1 |
| Antibodies |  |  |
| anti-pSTAT1 (Tyr701) | Cell Signaling Technology | 9167s |
| anti-pSTAT2 (Tyr690) | Cell Signaling Technology | 88410s |
| anti-STAT1 (p84/p91) | Santa Cruz Biotechnology Inc. | sc-346 |
| anti-STAT2 (A-7) | Santa Cruz Biotechnology Inc. | sc-1668 |
| anti-pSTAT3 (Tyr705) | Santa Cruz Biotechnology Inc. | sc-7993-r |
| anti-pSTAT3 (Tyr705) | Cell Signaling Technology | 9145s |
| anti-STAT4 (C46B10) | Cell Signaling Technology | 2653s |
| anti-STAT5A (4H1) | Cell Signaling Technology | 4807s |
| anti-STAT5B | Cell Signaling Technology | 34662s |
| anti-STAT6 | Cell Signaling Technology | 9362s |
| anti-Tubulin | Sigma-Aldrich | T9026 |
| anti-gH2AX | Cell Signaling Technology | 2577 |
| STING Pathway Antibody kit | Cell Signaling Technology | 38866 |
| Alexa Fluor® 647 Mouse Anti-Stat1 | BD Transduction Laboratories | 612597 |
| pStat2 Rabbit mAb (PE Conjugate) | Cell Signaling Technology | 77366s |
| HRP-conjugated anti-mouse antibody | Jackson ImmunoResearch | 115035146 |
| HRP-conjugated anti-rabbit antibody | Jackson ImmunoResearch | 111035144 |
| anti-Paxillin | BD Transduction Laboratories | 610052 |
| anti-Vinculin | Sigma-Aldrich | V9131 |
| anti-DM1A | Sigma-Aldrich | CP06 |
| Cy3 Donkey Anti-Rat IgG (H+L) | Jackson ImmunoResearch | 712-165-153 |
| Anti-mouse IgG 488 Conjugate | Cell Signaling Technology | 4408 |
| Deposited Data |  |  |
| Bulk MAR-seq data | This paper | GSE285030 |
| Recombinant DNA |  |  |
| pSpCas9(BB)-2A-Puro (PX459) V2.0 | Addgene | 62988 |
| pLPCX mito Grx1-roGFP2 | Addgene | 64977 |
| pLentiCRISPRv2 sgIRF3#5 | Addgene | 127642 |
| pLentiCRISPRv2-STING_gRNA3 | Addgene | 127640 |
| pLCRISPR-CMV-cGAS3 | Addgene | 102610 |
| pLenti6.3/V5-DEST-GFP | Addgene | 40125 |
| psPAX2 | Addgene | 12260 |
| pMD2.G | Addgene | 12259 |
| pLenti CMV-V5-GFP-DEST | This paper | N/A |
| PX459-hSTAT5A sg2 | This paper | N/A |
| PX459-hSTAT5B sg2 | This paper | N/A |
| PX459-hSTAT6 sg2 | This paper | N/A |
| pLenti CMV-V5-GFP-DEST | This paper | N/A |
| pLenti-CMV-STAT5A-Puro | This paper | N/A |
| pLPCX mito Grx1-roGFP2 | Addgene | 64977 |
| Oligonucleotides |  |  |
| Primers | This paper | Table S2 |
| Software and Algorithms |  |  |
| UTAP platform | ref<sup>62</sup> | N/A |
| Imaris 10.2 | Oxford Instruments | https://imaris.oxinst.com/ |
| Ingenuity Pathway Analysis | QIAGEN | https://shorturl.at/osoDQ |
| GraphpadPrism 10 | GraphPad Software Inc | https://www.graphpad.com/ |
| CytExpert Software | Beckman Coulter Inc. | https://shorturl.at/9e8UG |
| Flowjo | BD Biosciences | https://www.flowjo.com/ |
| LAS X Office | Leica Microsystems | https://shorturl.at/jW5eu |
| Integrative Genomics Viewer | Broad Institute | https://igv.org/ |
| ChIP-Atlas | ref<sup>72</sup> | https://chip-atlas.org/ |
| DepMap | ref<sup>25</sup> | https://depmap.org/portal |

**Table S2.**
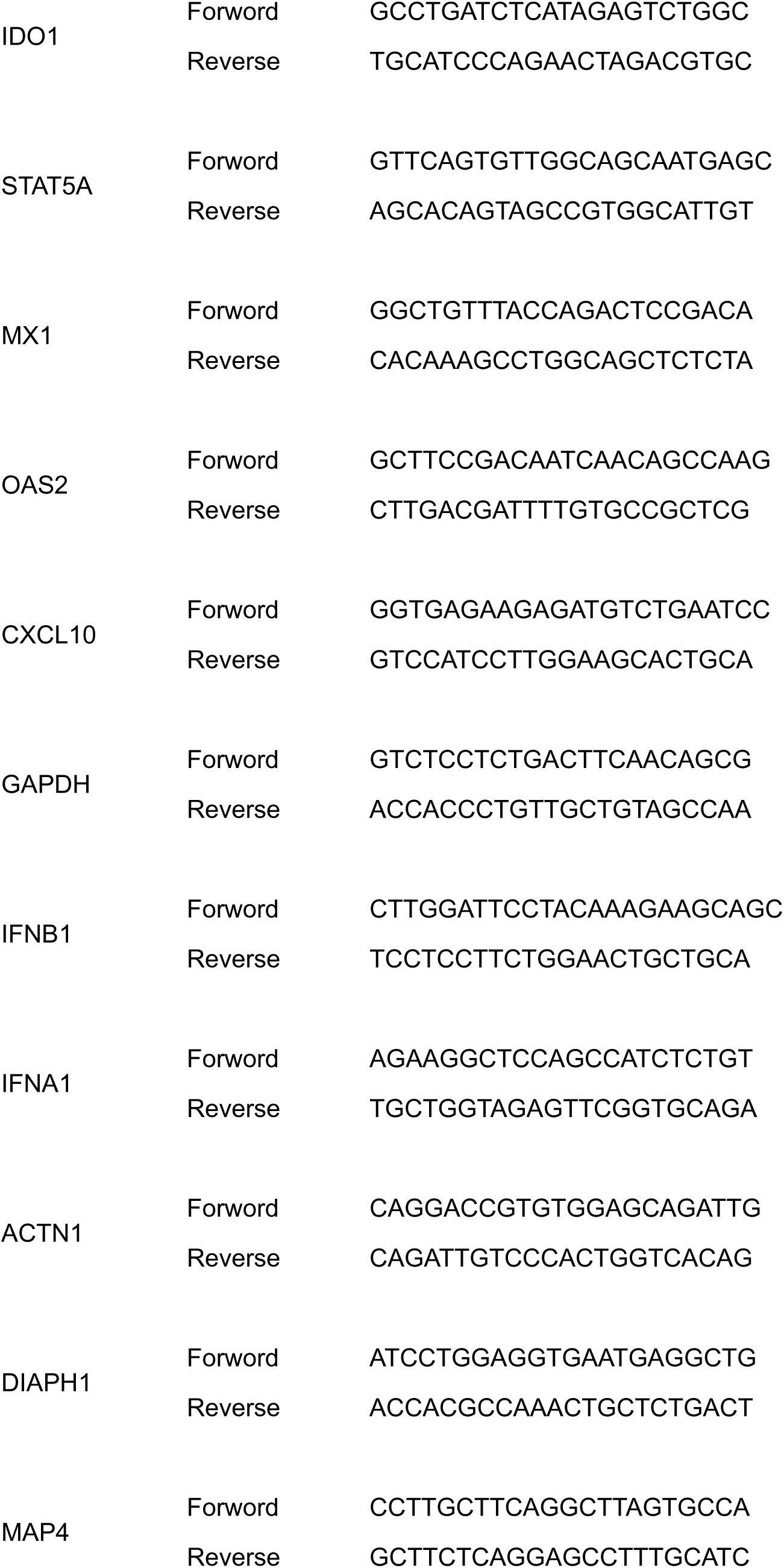
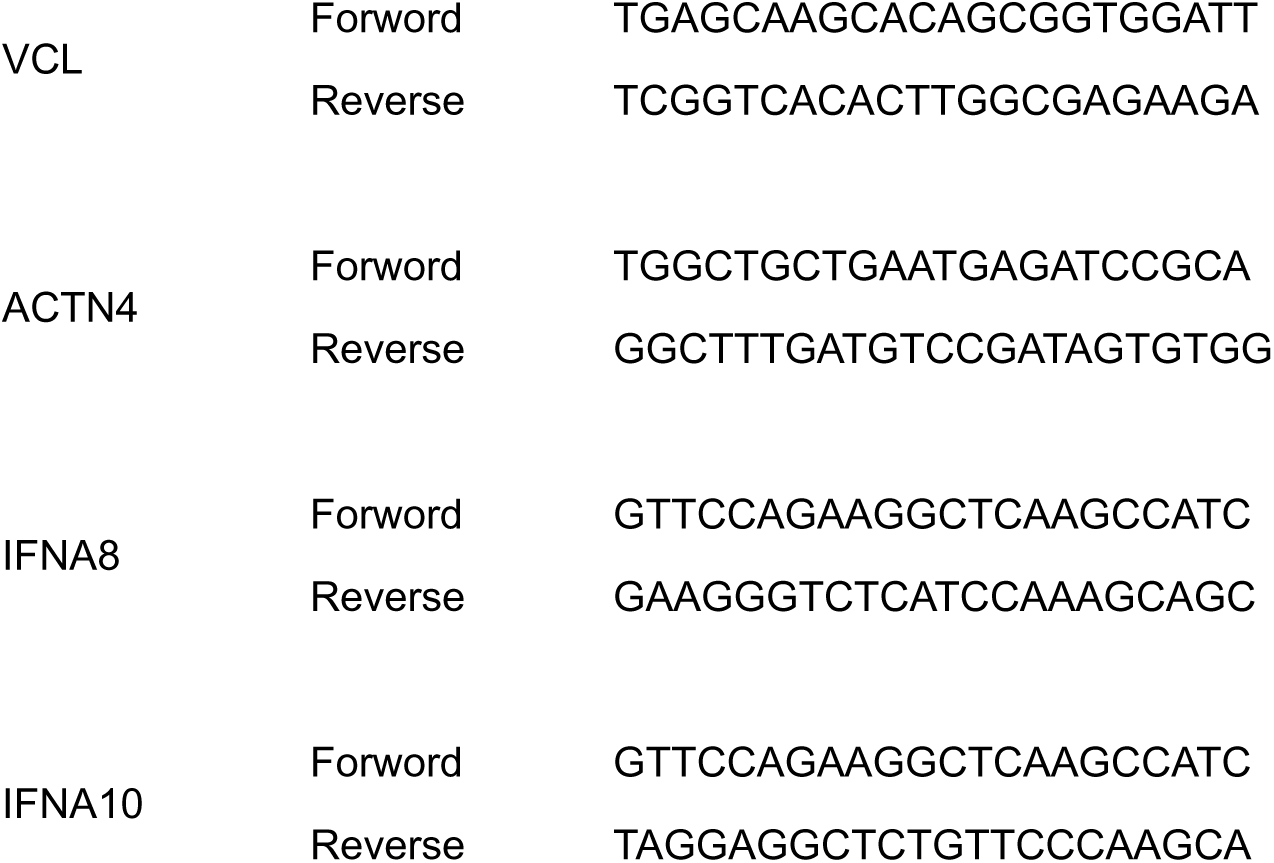
Primers for qPCR.

## References

1. Maurer, B., Kollmann, S., Pickem, J., Hoelbl-Kovacic, A. & Sexl, V. STAT5A and STAT5B—twins with different personalities in hematopoiesis and leukemia. Cancers (Basel) 11, 1726 (2019).

2. Rani, A. & Murphy, J. J. STAT5 in Cancer and Immunity. J. Interferon Cytokine Res. 36, 226– 237 (2016).

3. Monaghan, K. L. et al. Tetramerization of STAT5 promotes autoimmune-mediated neuroinflammation. Proc. Natl. Acad. Sci. U. S. A. 118, e2116256118 (2021).

4. Sheng, W. et al. STAT5 programs a distinct subset of GM-CSF-producing T helper cells that is essential for autoimmune neuroinflammation. Cell Res. 24, 1387–1402 (2014).

5. Verbsky, J. W. & Chatila, T. A. Chapter 23 - Immune Dysregulation Leading to Chronic Autoimmunity. in Stiehm’s Immune Deficiencies (eds. Sullivan, K. E. & Stiehm, E. R.) 497–516 (Academic Press, Amsterdam, 2014).

6. Hedl, M., Sun, R., Huang, C. & Abraham, C. STAT3 and STAT5 signaling thresholds determine distinct regulation for innate receptor–induced inflammatory cytokines, and *STAT3*/*STAT5* disease variants modulate these outcomes. J. Immunol. 203, 3325–3338 (2019).

7. Tolomeo, M., Meli, M. & Grimaudo, S. STAT5 and STAT5 inhibitors in hematological malignancies. Anticancer Agents Med. Chem. 19, 2036–2046 (2019).

8. Kaneshige, A. et al. A selective small-molecule STAT5 PROTAC degrader capable of achieving tumor regression in vivo. Nat. Chem. Biol. 19, 703–711 (2023).

9. Li, J., Tang, B., Miao, Y., Li, G. & Sun, Z. Targeting of STAT5 using the small molecule topotecan hydrochloride suppresses acute myeloid leukemia progression. Oncol. Rep. 50, 208–219 (2023).

10. Sehgal, P. B. Non-genomic STAT5-dependent effects at the endoplasmic reticulum and Golgi apparatus and STAT6-GFP in mitochondria. JAKSTAT 2, e24860 (2013).

11. Meier, J. A. & Larner, A. C. Toward a new STATe: The role of STATs in mitochondrial function. Semin. Immunol. 26, 20–28 (2014).

12. Casetti, L. et al. Differential contributions of STAT5A and STAT5B to stress protection and tyrosine kinase inhibitor resistance of chronic myeloid leukemia stem/progenitor cells. Cancer Res. 73, 2052–2058 (2013).

13. Bourgeais, J. et al. Oncogenic STAT5 signaling promotes oxidative stress in chronic myeloid leukemia cells by repressing antioxidant defenses. Oncotarget 8, 41876–41889 (2017).

14. Cholez, E. et al. Evidence for a protective role of the STAT5 transcription factor against oxidative stress in human leukemic pre-B cells. Leukemia 26, 2390–2397 (2012).

15. Illescas, M., Peñas, A., Arenas, J., Martín, M. A. & Ugalde, C. Regulation of mitochondrial function by the actin cytoskeleton. Front. Cell Dev. Biol. 9, 795838 (2021).

16. Moore, A. S., Wong, Y. C., Simpson, C. L. & Holzbaur, E. L. F. Dynamic actin cycling through mitochondrial subpopulations locally regulates the fission-fusion balance within mitochondrial networks. Nat. Commun. 7, 12886 (2016).

17. Hatch, A. L., Ji, W.-K., Merrill, R. A., Strack, S. & Higgs, H. N. Actin filaments as dynamic reservoirs for Drp1 recruitment. Mol. Biol. Cell 27, 3109–3121 (2016).

18. Fung, T. S., Chakrabarti, R. & Higgs, H. N. The multiple links between actin and mitochondria. Nat. Rev. Mol. Cell Biol. 24, 651–667 (2023).

19. Gourlay, C. W. & Ayscough, K. R. The actin cytoskeleton in ageing and apoptosis. FEMS Yeast Res. 5, 1193–1198 (2005).

20. Kim, J., Kim, H.-S. & Chung, J. H. Molecular mechanisms of mitochondrial DNA release and activation of the cGAS-STING pathway. Exp. Mol. Med. 55, 510–519 (2023).

21. Tzeng, H.-T., Chyuan, I.-T. & Lai, J.-H. Targeting the JAK-STAT pathway in autoimmune diseases and cancers: A focus on molecular mechanisms and therapeutic potential. Biochem. Pharmacol. 193, 114760 (2021).

22. Pollard, T. D. & Cooper, J. A. Actin, a central player in cell shape and movement. Science 326, 1208–1212 (2009).

23. Blanchoin, L., Boujemaa-Paterski, R., Sykes, C. & Plastino, J. Actin dynamics, architecture, and mechanics in cell motility. Physiol. Rev. 94, 235–263 (2014).

24. DepMap, B. DepMap 24Q2 Public. Figshare+ 10.25452/FIGSHARE.PLUS.25880521.V1 (2024).

25. Tsherniak, A. et al. Defining a cancer dependency map. Cell 170, 564–576.e16 (2017).

26. Hollenbeck, P. J. & Saxton, W. M. The axonal transport of mitochondria. J. Cell Sci. 118, 5411–5419 (2005).

27. Moore, A. S. & Holzbaur, E. L. F. Mitochondrial-cytoskeletal interactions: dynamic associations that facilitate network function and remodeling. Curr. Opin. Physiol. 3, 94–100 (2018).

28. Yadav, T., Gau, D. & Roy, P. Mitochondria-actin cytoskeleton crosstalk in cell migration. J. Cell. Physiol. 237, 2387–2403 (2022).

29. Kim, H. J. et al. Dynamin-related protein 1 controls the migration and neuronal differentiation of subventricular zone-derived neural progenitor cells. Sci. Rep. 5, 15962 (2015).

30. Ji, W.-K., Hatch, A. L., Merrill, R. A., Strack, S. & Higgs, H. N. Actin filaments target the oligomeric maturation of the dynamin GTPase Drp1 to mitochondrial fission sites. Elife 4, e11553 (2015).

31. De Vos, K. J., Allan, V. J., Grierson, A. J. & Sheetz, M. P. Mitochondrial function and actin regulate dynamin-related protein 1-dependent mitochondrial fission. Curr. Biol. 15, 678–683 (2005).

32. Rogakou, E. P., Pilch, D. R., Orr, A. H., Ivanova, V. S. & Bonner, W. M. DNA double-stranded breaks induce histone H2AX phosphorylation on serine 139. J. Biol. Chem. 273, 5858–5868 (1998).

33. Holmström, K. M. & Finkel, T. Cellular mechanisms and physiological consequences of redox-dependent signalling. Nat. Rev. Mol. Cell Biol. 15, 411–421 (2014).

34. Perry, S. W., Norman, J. P., Barbieri, J., Brown, E. B. & Gelbard, H. A. Mitochondrial membrane potential probes and the proton gradient: A practical usage guide. Biotechniques 50, 98– 115 (2011).

35. Kasozi, D., Mohring, F., Rahlfs, S., Meyer, A. J. & Becker, K. Real-time imaging of the intracellular glutathione redox potential in the malaria parasite Plasmodium falciparum. PLoS Pathog. 9, e1003782 (2013).

36. Geissel, F., Lang, L., Husemann, B., Morgan, B. & Deponte, M. Deciphering the mechanism of glutaredoxin-catalyzed roGFP2 redox sensing reveals a ternary complex with glutathione for protein disulfide reduction. Nat. Commun. 15, 1733 (2024).

37. Volkman, H. E., Cambier, S., Gray, E. E. & Stetson, D. B. Tight nuclear tethering of cGAS is essential for preventing autoreactivity. Elife 8, e47491 (2019).

38. Samson, N. & Ablasser, A. The cGAS–STING pathway and cancer. Nature Cancer 3, 1452– 1463 (2022).

39. Cai, X., Chiu, Y.-H. & Chen, Z. J. The cGAS-cGAMP-STING pathway of cytosolic DNA sensing and signaling. Mol. Cell 54, 289–296 (2014).

40. Zhang, L. et al. Mitochondrial STAT5A promotes metabolic remodeling and the Warburg effect by inactivating the pyruvate dehydrogenase complex. Cell Death Dis. 12, 634 (2021).

41. Gourlay, C. W. & Ayscough, K. R. The actin cytoskeleton: a key regulator of apoptosis and ageing? Nat. Rev. Mol. Cell Biol. 6, 583–589 (2005).

42. Ligon, L. A. & Steward, O. Role of microtubules and actin filaments in the movement of mitochondria in the axons and dendrites of cultured hippocampal neurons. J. Comp. Neurol. 427, 351– 361 (2000).

43. Cho, M. J., Kim, Y. J., Yu, W. D., Kim, Y. S. & Lee, J. H. Microtubule integrity is associated with the functional activity of mitochondria in HEK293. Cells 10, 3600 (2021).

44. Campello, S. & Scorrano, L. Mitochondrial shape changes: orchestrating cell pathophysiology. EMBO Rep. 11, 678–684 (2010).

45. Korobova, F., Ramabhadran, V. & Higgs, H. N. An actin-dependent step in mitochondrial fission mediated by the ER-associated formin INF2. Science 339, 464–467 (2013).

46. Gatti, P., Schiavon, C., Cicero, J., Manor, U. & Germain, M. Mitochondria- and ER-associated actin are required for mitochondrial fusion. Nat. Commun. 16, 451 (2025).

47. Trono, P., Tocci, A., Musella, M., Sistigu, A. & Nisticò, P. Actin Cytoskeleton Dynamics and Type I IFN-Mediated Immune Response: A Dangerous Liaison in Cancer. Biology 10, 913 (2021).

48. Seo, D. & Gammon, D. B. Manipulation of the host cytoskeleton by viruses: Insights and mechanisms. Viruses 14, 1586 (2022).

49. Kloc, M., Uosef, A., Wosik, J., Kubiak, J. Z. & Ghobrial, R. M. Virus interactions with the actin cytoskeleton-what we know and do not know about SARS-CoV-2. Arch. Virol. 167, 737–749 (2022).

50. Colonne, P. M., Winchell, C. G. & Voth, D. E. Hijacking host cell highways: Manipulation of the host actin cytoskeleton by obligate intracellular bacterial pathogens. Front. Cell. Infect. Microbiol. 6, (2016).

51. Lamason, R. L. & Welch, M. D. Actin-based motility and cell-to-cell spread of bacterial pathogens. Curr. Opin. Microbiol. 35, 48–57 (2017).

52. Igelmann, S., Neubauer, H. A. & Ferbeyre, G. STAT3 and STAT5 Activation in Solid Cancers. Cancers 11, 1428–1437 (2019).

53. Halim, C. E., Deng, S., Ong, M. S. & Yap, C. T. Involvement of STAT5 in Oncogenesis. Biomedicines 8, 316–336 (2020).

54. Li, M. et al. Roles of the cytoskeleton in human diseases. Mol. Biol. Rep. 50, 2847–2856 (2023).

55. Schwarz, T. L. Mitochondrial trafficking in neurons. Cold Spring Harb. Perspect. Biol. 5, a011304–a011304 (2013).

56. Xia, T., Konno, H. & Barber, G. N. Recurrent loss of STING signaling in melanoma correlates with susceptibility to viral oncolysis. Cancer Res. 76, 6747–6759 (2016).

57. Kim, S. K. & Cho, S. W. The evasion mechanisms of cancer immunity and drug intervention in the tumor microenvironment. Front. Pharmacol. 13, 868695 (2022).

58. Hoffmann, L. et al. Cofilin1 oxidation links oxidative distress to mitochondrial demise and neuronal cell death. Cell Death Dis. 12, 953 (2021).

59. Hoffmann, L., Rust, M. B. & Culmsee, C. Actin(g) on mitochondria - a role for cofilin1 in neuronal cell death pathways. Biol. Chem. 400, 1089–1097 (2019).

60. Ran, F. A. et al. Genome engineering using the CRISPR-Cas9 system. Nat. Protoc. 8, 2281– 2308 (2013).

61. Shemesh, M., Lochte, S., Piehler, J. & Schreiber, G. IFNAR1 and IFNAR2 play distinct roles in initiating type I interferon-induced JAK-STAT signaling and activating STATs. Sci. Signal. 14, eabe4627 (2021).

62. Kohen, R. et al. UTAP: User-friendly Transcriptome Analysis Pipeline. BMC Bioinformatics 20, 154–161 (2019).

63. Martin, M. Cutadapt removes adapter sequences from high-throughput sequencing reads. EMBnet J. 17, 10 (2011).

64. Dobin, A. et al. STAR: ultrafast universal RNA-seq aligner. Bioinformatics 29, 15–21 (2013).

65. Anders, S., Pyl, P. T. & Huber, W. HTSeq--a Python framework to work with high-throughput sequencing data. Bioinformatics 31, 166–169 (2015).

66. Love, M. I., Huber, W. & Anders, S. Moderated estimation of fold change and dispersion for RNA-seq data with DESeq2. Genome Biol. 15, 550 (2014).

67. Benjamini, Y. & Hochberg, Y. Controlling the false discovery rate: A practical and powerful approach to multiple testing. J. R. Stat. Soc. Series B Stat. Methodol. 57, 289–300 (1995).

68. Aller, I., Rouhier, N. & Meyer, A. J. Development of roGFP2-derived redox probes for measurement of the glutathione redox potential in the cytosol of severely glutathione-deficient rml1 seedlings. Front. Plant Sci. 4, 506 (2013).

69. Zapatero-Solana, E. et al. Oxidative stress and mitochondrial dysfunction in Kindler syndrome. Orphanet J. Rare Dis. 9, 211 (2014).

70. Zoler, E. et al. Promiscuous Janus kinase binding to cytokine receptors modulates signaling efficiencies and contributes to cytokine pleiotropy. Sci. Signal. 17, eadl1892 (2024).

71. Urin, V., Shemesh, M. & Schreiber, G. CRISPR/Cas9-based Knockout Strategy Elucidates Components Essential for Type 1 Interferon Signaling in Human HeLa Cells. J. Mol. Biol. 431, 3324– 3338 (2019).

72. Zou, Z., Ohta, T. & Oki, S. ChIP-Atlas 3.0: a data-mining suite to explore chromosome architecture together with large-scale regulome data. Nucleic Acids Res. 52, W45–W53 (2024).

